# NtcA, LexA and heptamer repeats involved in the multifaceted regulation of DNA repair genes *recF, recO* and *recR* in the cyanobacterium *Nostoc* PCC7120

**DOI:** 10.1101/2022.06.06.494920

**Authors:** Mitali Pradhan, Arvind Kumar, Anurag Kirti, Sarita Pandey, Hema Rajaram

## Abstract

Regulation of DNA repair genes in cyanobacteria is an unexplored field despite some of them exhibit high radio-resistance. With the RecF pathway speculated to be the major double strand break repair pathway in *Nostoc* sp. strain PCC7120, the regulation of *recF, recO* and *recR* genes was investigated. Bioinformatic approach-based identification of promoter and regulatory elements was validated using qRT-PCR analysis, reporter gene and DNA binding assays. Different deletion constructs of the upstream regulatory regions of these genes were analysed in host *Nostoc* as well as heterologous system *Escherichia coli*. Studies revealed: (1) Positive regulation of all three genes by NtcA, (2) Negative regulation by LexA, (3) Involvement of contiguous heptamer repeats with/without its yet to be identified interacting partner in regulating (i) binding of NtcA and LexA to *recO* promoter and its translation, (ii) transcription or translation of *recF*, (4) Translational regulation of *recF* and *recO* through non-canonical and distant S.D. sequence and of *recR* through a rare initiation codon. Presence of NtcA either precludes binding of LexA to AnLexA-Box or negates its repressive action resulting in higher expression of these genes under nitrogen-fixing conditions in *Nostoc*. Thus, in *Nostoc*, expression of *recF, recO* and *recR* genes is intricately regulated through multiple regulatory elements/proteins. Contiguous heptamer repeats present across the *Nostoc* genome in the vicinity of start codon or promoter is likely to have a global regulatory role. This is the first report on the involvement of NtcA and heptamer repeats in regulation of DNA repair genes in any organism.

## 1. INTRODUCTION

Cyanobacteria have been exposed to various environmental stresses including γ-radiation during it’s over 3 billion years of existence on Earth (1). Some of the cyanobacterial species exhibit high radio-resistance, e.g. *Chroococcidiopsis* (2), *Arthrospira* sp, PCC8005 (3) and *Nostoc* (*Anabaena*) sp. PCC7120 (4). The ability of *Nostoc* sp. PCC7120 to re-stitch its DNA during post-irradiation recovery (4) has been speculated to be through the RecF pathway of double strand break (DSB) repair pathways (5). In fact, in the highly radioresistant *Deinococcus radiodurans* which lacks the RecBC proteins, the RecF pathway is the predominant double strand break (DSB) repair pathway, both in HR and extended synthesis dependent strand annealing (ESDSA) (6,7). RecF pathway (RecFOR/RecOR) has been extensively studied in *Escherichia coli*, and involves several proteins *viz*. RecF, RecO, RecR, RecJ, RecN, RecQ, RuvA, RuvB, RuvC and SSB proteins (8,9).

Regulation of expression of the *recF,O,R* genes is essential given its important role in DNA repair and recombination, and has been majorly and extensively studied in *Escherichia coli*. One of the well-established regulation of these genes is through the SOS-response, wherein they are negatively regulated by the repressor protein LexA (10), which binds to the LexA (SOS)-Box [CTG(N_8_)CAG] (11). In *E. coli*, a hexapyrimidine tract located just upstream of the ribosome interactive sequence (epsilon) was shown to be involved in its regulation at the translational level through mutational analysis (12). In *E. coli*, the *recF* gene lies in a cluster of DNA synthesis genes *dnaA-dnaN-recF-gyrB*, and has three overlapping promoters located well within the *dnaN* gene, with two of them capable of initiating transcription towards the *recF* gene, while the 3^rd^ does so in the opposite direction (13). The predominant mRNA is the *dnaA-dnaN-recF* mRNA which has three tandem transcriptional terminators upstream of the *recF* gene, and is likely to be involved in the down-regulation of the *recF* gene (14). The transcription of these genes is dependent on the growth phase of the cells, with *dnaN* and *recF* expressed predominantly from the *dnaA* promoters in the exponential phase, while under stationary phase, their transcription is high and independent of *dnaA*, but dependent on RpoS (15). In *Azotobacter vinelandii*, the UV-inducible *recF* gene has been speculated to be regulated by LexA, based on the presence of a LexA-box upstream of the *recF* gene (16). In *E. coli* and *Salmonella typhimurium, recO* is a part of *rnc-era-recO* operon, which is an autoregulated operon with Ribonuclease III (RNase III) coded by *rnc* gene cleaving the stem-loop structure in 5’ untranslated leader sequence, thus initiating rapid decay of *m*RNA and lower transcript levels (17,18). The presence of *recO* as part of the *rnc-era-recO* operon is well conserved and also observed in *Pseudomonas aeruginosa* and *Coxiella burnetti* (19,20). In *A. actinomycetemcomitans, recO* was found to be regulated by FurA (21). In the radioresistant bacteria *D. radiodurans*, the *recF, recO* and *recR* genes are speculated to be under the regulation of G4 quadruplex found in their promoter region (22). The *recR* gene of *E. coli* is part of the *dnaX*-*orf12*-*recR* operon (23), with the translation of *recR* found to be coupled with that of *orf12* (24).

Cyanobacteria has several global transcriptional regulators of which NtcA (25), LexA (26,27) and PipX, a co-activator of NtcA (28) have been shown to function both as activators and repressors. NtcA a well-known nitrogen response regulator in cyanobacteria was initially identified as binding to the consensus sequence GTA-N_8_-TAC near the promoter region (25), and 158 transcriptional start sites as potential targets for regulation by NtcA were identified subsequently (29). Using the high-throughput techniques of ChIP-Seq and RNA-Seq, 51 up-regulated genes and 28 down-regulated genes belonging to NtcA direct regulon was unearthed in *Synechocystis* PCC6803, with the binding site defined as GT-N_10_-AC (30). In *Anabaena* PCC7120, ChIP Seq analysis revealed 2424 putative DNA regions of which 865 were in the promoter regions corresponding to a wide NtcA binding site defined mostly as GT-N_10_-AC, but also included a few wherein either only the left arm (GTA) or the right arm (TAC) is conserved, provided the end bases are invariant G and C and spacing between ‘G’ and ‘C’ is 12 (31). The well-known SOS-response regulator LexA transcriptionally regulates several genes beyond those involved in DNA repair through binding to the AnLexA-Box [AGT-N_4-11_-ACT] allowing for one base mismatch in case of a 4-base palindrome (27,32). In fact, in the unicellular cyanobacteria, *Synechocystis* PCC6803, LexA has been shown to regulate genes involved in C-metabolism but not those in DNA repair (33,34).

The *recF, recO* and *recR* genes in the cyanobacterium, *Nostoc* sp. strain PCC 7120, henceforth referred to as *Nostoc* 7120, exhibited differential expression at transcript and protein levels, with *recF* expressed least at transcript level and maximal at the protein level (35), suggesting the possibility of multilevel regulation of these genes. Using the heterologous system i.e. *E. coli* and the host strain *Nostoc* 7120, the role of different regulatory elements (AnLexA-Box, NtcA-Box and heptamer repeats) in controlling the expression of *recF, recO* and *recR* genes has been evaluated in this paper. This is the first instance wherein the role of NtcA in regulating DNA repair genes has been reported in any cyanobacteria. Additionally, the possibility of contiguous heptamer repeats having a regulatory role has also been discussed along with promoter activity and transcription profile of these genes during post-irradiation recovery.

## 2. MATERIALS AND METHODS

### 2.1 Organism, growth conditions and γ-radiation stress

*E. coli* cells were grown in Luria-Bertani (LB) medium in broth with shaking (150 rpm) or on 1.5 % LB agar plates at 37 °C. Antibiotics [50 µg Kanamycin mL^-1^ (Kan_50_), 34 µg chloramphenicol mL^-1^ (Cm_34_) or 100 µg carbenicillin mL^-1^ (Cb_100_)] were added into the medium when required. *Nostoc* 7120 and recombinant *Nostoc* strains were grown axenically in BG-11 liquid medium, pH 7.0 with (BG11, N^+^) or without (BG11, N^-^)17 mM NaNO_3_ (36), under stationary conditions with continuous illumination (30 µE m^-2^ s^-1^) at 27 ± 2 °C. Additionally, the antibiotic neomycin (Neo) was added (at 12.5 μg mL^-1^ in liquid medium or 25 μgmL^-1^ in solid medium) for recombinant *Nostoc* strains. Three-day-old *Nostoc* cultures were concentrated to 10 µg mL^-1^ Chlorophyll (Chl) *a* density and subjected to 6 kGy ^60^Co γ-irradiation and post-irradiation recovery (PIR) as described earlier (37). Growth was measure in terms of Chl *a* density as described earlier (38).

### 2.2 Cloning and overexpression of *Nostoc* 7120 NtcA in *E. coli*

The *ntcA* gene of *Nostoc* 7120 was PCR amplified from genomic DNA using the gene specific primers (Table S1A), digested with *Nde*I and *Bam*HI and cloned in pet16b at the same restriction site to generate the recombinant plasmid pET*ntcA* (Table S2). Overexpression of NtcA was achieved in *E. coli* BL21(*plysS*) DE3 cells bearing this plasmid (Table S3) induced with 1mM IPTG for 18 h at 20 °C.

### 2.3 Cloning of promoter fragments and construction of recombinant *E. coli* and *Nostoc* strains

PCR amplification was carried out using from genomic DNA of *Nostoc* 7120 using 1 µmole each specific forward and reverse primers (Table S1A), 100 µM dNTPs, and Taq DNA Polymerase in the Taq Buffer provided with kit (BRIT Vashi) to generate the promoter fragments indicated in Table S1B. The different amplicons were cloned in pAM1956 (39) to yield the plasmid constructs pAM-P_*abc*_ (*abc* referring to different amplicons) (Table S2) and maintained in *E. coli* DH5α and HB101. These plasmids were (i) co-transformed with either pET16b, pET*lexA* or pET*ntcA* into *E. coli* BL21(p*lysS*) cells, and (ii) conjugated into *Nostoc* 7120 as described earlier (40) to generate different recombinant strains as listed in Table S3. DNA inserts of all the recombinant plasmids used were sequenced on both the strands and compared with the available sequence in data base (https://www.genome.jp/kegg-bin/show_organism?org=ana) to confirm that no error had been incorporated during PCR amplification.

### 2.4 Measurement of promoter activity

For assessing promoter activity in *E. coli*, mid-logarithmic phase cultures were further incubated in the absence or presence of 1 mM IPTG and Green Fluorescent Protein (GFP) expression measured at 0 h and 2 h in case of LexA-overexpressing cells and at the end of 18 h for NtcA-overexpressing cells as described earlier (27). For *Nostoc* cultures, expression was measured for 3-day-old cultures and at specified time points of PIR and corresponding controls as described earlier (41). Fluorescence of samples was measured using JASCO FP6500 spectrofluorometer (λ_ex_: 489 nm, λ_em_: 509 nm). The data was represented as fluorescence in Arbitrary Units per OD_750_ for *E. coli* and µg Chl *a* for *Nostoc* cultures. All experiments were carried out with 3 sets of technical replicates and 2 sets of biological replicates.

### 2.5 RNA isolation, cDNA synthesis and Real time PCR

Total RNA was extracted from 3-day old culture of *Nostoc* cultures subjected to different growth conditions, quantified and purity checked in terms of absence of DNA as contaminant as described earlier (27). For cDNA synthesis, 1 μg of total RNA was used to reverse-transcribe in a total 20 μL reaction volume, using ReadyScript cDNA Synthesis Mix (Sigma, USA) as per manufacturer’s protocol. Gene specific primers (Table 1C) designed using NCBI Primer-Blast to generate 100-150 bp amplicons were used for amplification. qPCR was carried out with using Realtime PCR machine (Rotor-□Gene Q, Qaigen, Germany) using 2X FastStart SYBR Green Master (Roche, Germany). In brief, 10 ng equivalent cDNA along with primers and other reagents was carried out in 20 µL reaction volume and cycling condition as per manufacturer’s protocol with varying annealing temperature. 16S rDNA was taken as reference gene to normalize the transcript levels of the genes tested and the fold differences of each sample was calculated using the 2^-ΔΔCt^ method (42). The threshold cycle (Ct) was automatically determined for each reaction using the system set with default parameters. The specificity of PCR was determined using the melt curve analysis of amplified products.

### 2.6 Electrophoretic Mobility Shift Assay (EMSA)

EMSA experiments of the different DNA fragments with varying concentrations of purified *Nostoc* LexA protein was carried out as described earlier (27). The densitometric analysis of the images was carried out suing ImageJ software (43), and the graph between % unbound DNA and LexA concentration used to determine the binding affinity as described earlier (27).

### 2.7 Statistical Analysis

A minimum of two biological replicates and three technical replicates were used for performing all experiments. Values are represented as mean ± Standard deviation (n=3) and statistical significance was determined using t-test and represented as * (p<0.05). ** (p<0.01) and *** (p<0.001).

## 3. RESULTS AND DISCUSSION

### 3.1 Expression of *recF, recO* and *recR* in response to LexA overexpression and N-status in *Nostoc* 7120

Having predicted the presence of AnLexA-Box upstream of *recF, recO* and *recR* genes earlier (5), the regulation of these genes by LexA was validated by comparing their transcript levels in vector control, AnpAM (44) and LexA-overexpressing An*lexA*^+^ (27) cells. Transcript levels of *recF, recO* and *recR* were 2-fold, 2.22-fold and 3.12-fold lower in An*lexA*^+^ compared to AnpAM cells respectively (Fig. 1A), thereby confirming negative regulation of these genes by LexA. Additionally, comparison of the transcript levels under nitrogen-fixing (N-fix) and N-supplemented (N-sup) conditions in AnpAM revealed 2.47-fold, 2.42-fold and 3.07-fold higher transcript levels of *recF, recO* and *recR* respectively under N-fix conditions compared to N-sup conditions (Fig. 1A). This suggested N-status-dependent regulation of these three DNA repair genes as well.

**Figure 1.**
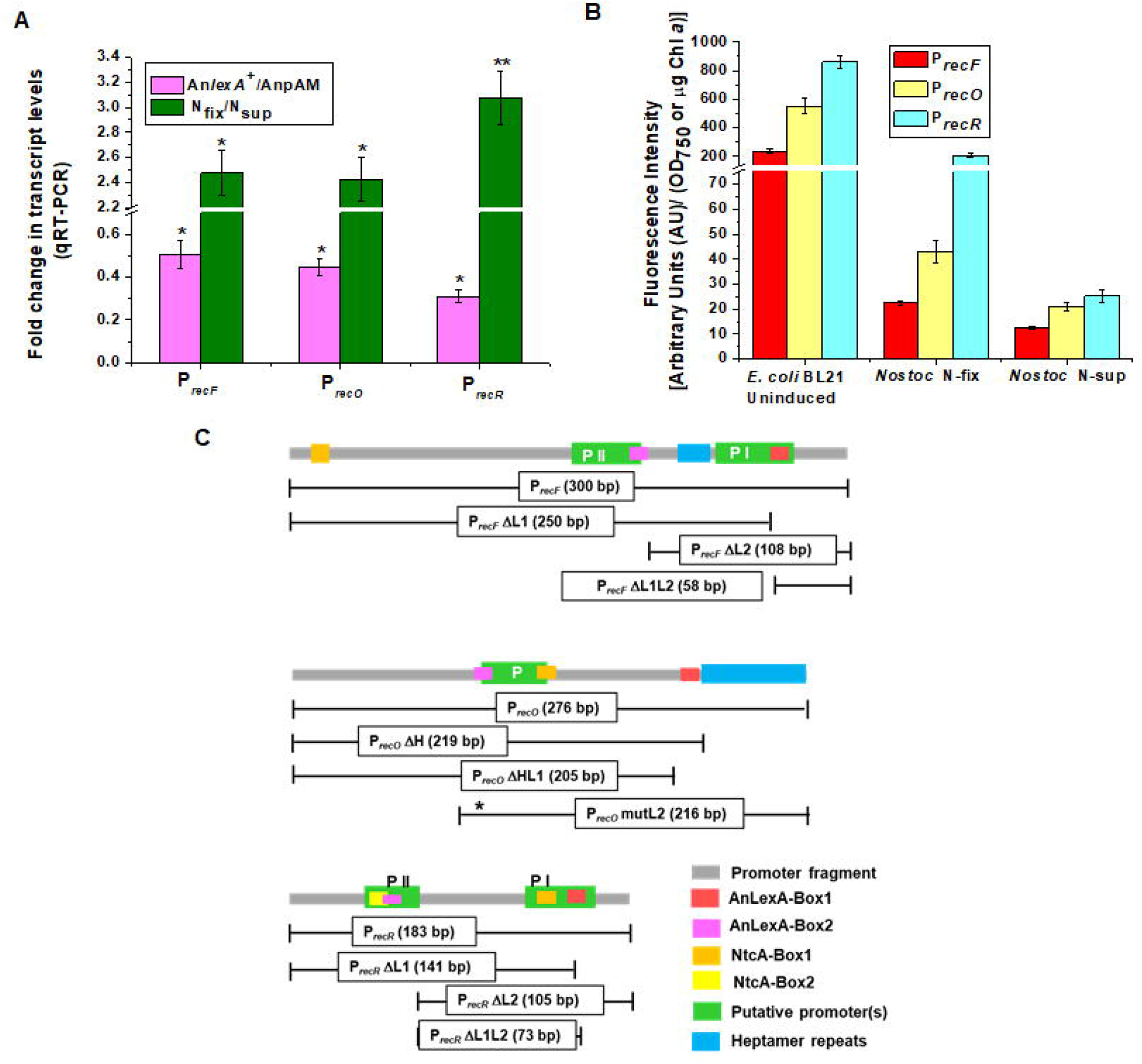
Expression and regulatory regions of *recF, recO* and *recR* genes of *Nostoc* PCC7120. (A) Fold change in the transcript levels of *recF, recO* and *recR* in 3-day-old (i) An*lexA*^+^ (LexA overexpressing *Nostoc* strain) compared to that in vector control AnpAM, and (ii) AnpAM cultures grown under nitrogen-fixing (N-fix) conditions compared to that under nitrogen-supplemented (N-sup) conditions was analysed by qRT-PCR. Values are represented as mean ± Standard deviation (n=3) and statistically significant differences indicated as * (p<0.05). ** (p<0.01) and *** (p<0.001). (B) Expression of *recF, recO* and *recR* promoters in *E. coli* BL21 (*plysS*) cells under uninduced conditions and in 3-day-old *Nostoc* cultures under N-fix and N-sup conditions. Promoter activity was measured as GFP fluorescence (λ_ex_ 489 nm, λ_em_ 509 nm) (arbitrary units, AU) and expressed per unit O.D._750_ for *E. coli* and per µg Chl *a* content for *Nostoc* cultures. (C) Schematic representation of the DNA fragments corresponding to the upstream region of the *recF/O/R* genes used for promoter analysis. The predicted -10 promoter regions, AnLexA-boxes, NtcA Box and heptamer repeats are shown as coloured rectangles. The regions corresponding to different constructs used in this study are also shown as line diagram with their size.

To assess this further, a ∼300 bp region upstream of these genes were analysed for promoter activity using GFP reporter assay under different conditions in *E. coli* as well as *Nostoc* 7120. The different plasmid constructs pAM-P_*recF*_, pAM-P_*recO*_ and pAM-P_*recR*_ generated using the specific primer sets (Tables S1-3) exhibited GFP expression both in *E. coli* and *Nostoc* cells, level of expression in *E. coli* being 5-10-fold higher (Fig. 1B). The higher promoter activity under N-fix conditions compared to N-sup (1.79-fold, 2.04-fold and 8.19-fold for P_*recF*_, P_*recO*_ and P_*recR*_ respectively) (Fig. 1B) correlated with the trend observed for the corresponding transcript levels (Fig. 1A). This suggested a possible direct or indirect regulation through NtcA. In both *E. coli* and *Nostoc*, the promoter strength was found to be in the order P_*recF*_ < P_*recO*_ < P_*recR*_ (Fig. 1B).

### 3.2 Identification of the regulatory elements upstream of the *recF, O, R* genes

Sequence analysis of P_*recF*_, P_*recO*_ and P_*recR*_ fragments revealed the presence of (i) two promoter elements, predicted using the BPROM software (http://www.softberry.com/berry, 45), designated as PI and PII in P_*recF*_ and P_*recR*_ and one (P) in P_*recO*_, (ii) two AnLexA-boxes based on the earlier defined consensus sequence (27,32), AnLexA-Box1 and AnLexA-Box2 in all three, with AnLexA-Box1 corresponding to the one identified earlier (5), (iii) NtcA-like boxes, based on the earlier defined consensus sequence (31), two upstream of *recR* and one each upstream of *recF* and *recO*, and (iv) two distinct heptamer repeats in P_*recF*_ and P_*recO*_ (Figs. 1C, S1). Different deletion constructs of the promoters were made lacking one or both the AnLexA-boxes as shown in Fig. 1C. Binding studies with purified *Nostoc* LexA revealed that presence of either of the AnLexA-boxes was sufficient for *in vitro* binding for all three promoters, while no binding was observed when both the AnLexA-boxes were deleted (Fig. 2). The binding affinity calculated graphically based on the amount of unbound DNA as shown in Fig. S2 was found to be in the range of 70 to 110 nM irrespective of the presence of one or two AnLexA-boxes (Table S4), which correlated well with the binding affinity of LexA determined earlier (27).

**Figure 2.**
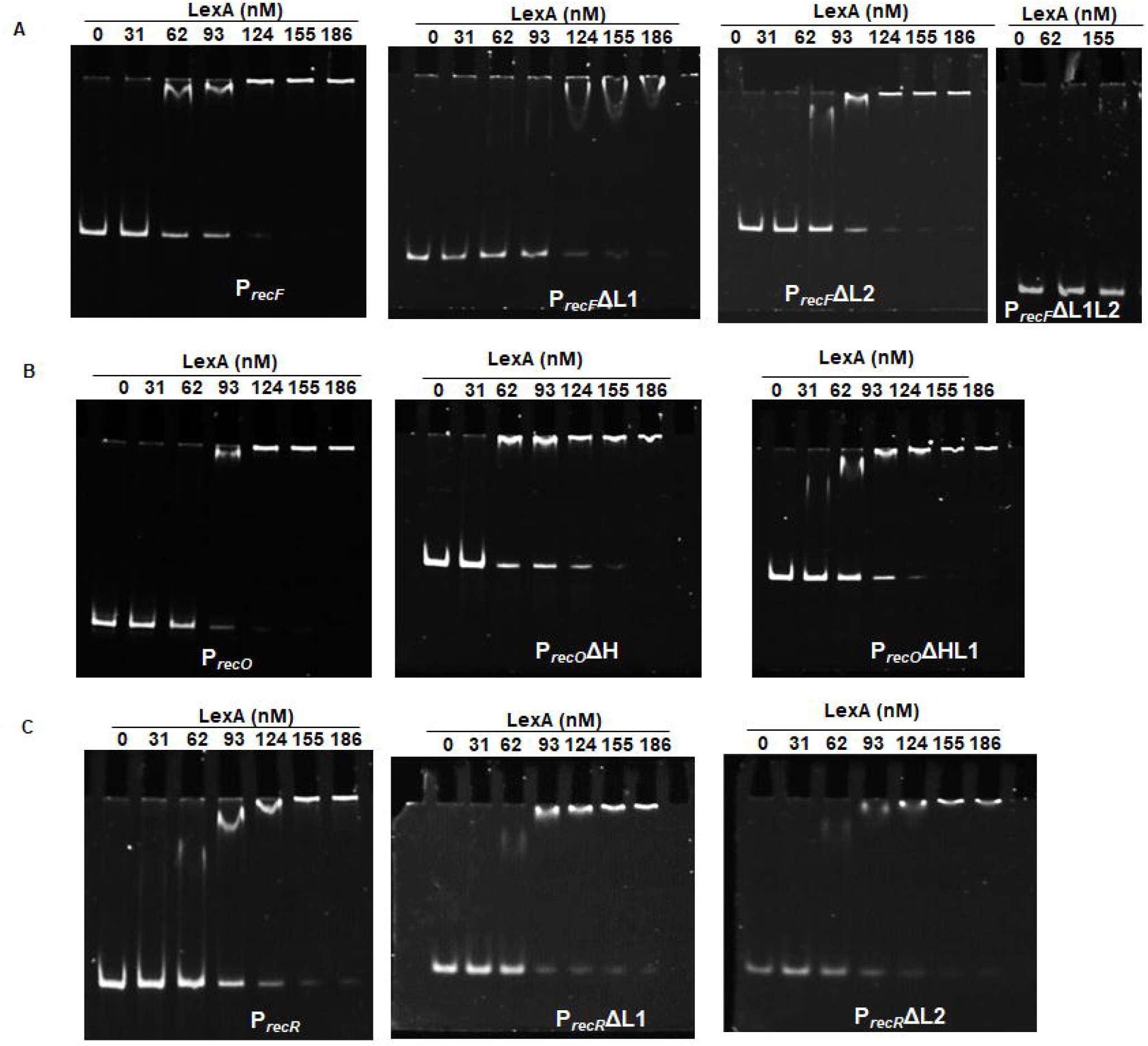
Binding of *Nostoc* LexA to different promoter constructs. Electrophoretic mobility shift assay was carried out using 25 ng of different DNA fragments with different concentrations of *Nostoc* LexA with (A) P_*recF*_, P_*recF*_ΔL1, P_*recF*_ΔL2 and P_*recF*_ΔL1L2, (B) P_*recO*_, P_*recO*_ΔH and P_*recO*_ΔHL1, and (C) P_*recR*_, P_*recR*_ΔL1 and P_*recR*_ΔL2. The lower band corresponds to free or unbound DNA and the band with retarded mobility to the bound DNA.

Shine-Delgarno (S.D.) motifs are weakly conserved in cyanobacteria, and the preferred anti-SD sequence has been predicted for unicellular cyanobacteria, *Synechocystis* PCC6803 and *Microcystis aeruginosa* (46), but not for any filamentous cyanobacteria. Hence, for predicting the probable SD sequence for the *recF, O, R* genes, the known sequence of 3’ end of mature 16S rRNA of *E. coli* (3’AUUCCUCCACUAGGUUGGCG---5’) (46) was used. The probable SD sequence for *recF, recO* and *recR* genes were identified as ‘ATCC’, ‘CCAA’ and ‘GGAG’ respectively, the corresponding distance from the start codon being 12, 6 and 11 bases respectively (Fig. S1).

In order to identify the intricacies of the regulation of *recF, recO* and *recR* genes of *Nostoc*, the full length and deletion promoter constructs were analysed for promoter activity in *E. coli* and *Nostoc*.

### 3.3 Regulation of expression of *recF* gene

In *E. coli*, all three constructs (P_*recF*_, P_*recF*_ΔL1 and P_*recF*_ΔL2) exhibited promoter activity (Figs. 1B, 3A) indicating that both PI and PII promoters are active. Deletion of both the promoters as in P_*recF*_ΔL1L2 (Fig. 1C) yielded no promoter activity (data not shown). Activity of P_*recF*_ΔL1 (having PII promoter) was similar to that of P_*recF*_ (Fig. 3A) suggesting that PII could be the preferred promoter for initiating transcription of *Nostoc recF* in *E. coli*. However, the activity of P_*recF*_ΔL2 (having PI promoter) was 1.6-fold higher than that of P_*recF*_ (Fig. 3A) indicating that the strength of PI is higher concurring with the bioinformatics analysis of the promoters (data not shown). In 3-day-old *Nostoc* cultures compared to P_*recF*_, activity of P_*recF*_ΔL1 was 1.6-fold higher under N-fix conditions, but similar under N-sup conditions, while the reverse was true for P_*recF*_ΔL2, wherein its activity was similar to that of P_*recF*_ under N-fix conditions, but 1.8-fold higher under N-sup conditions (Fig. 3A).

**Figure 3.**
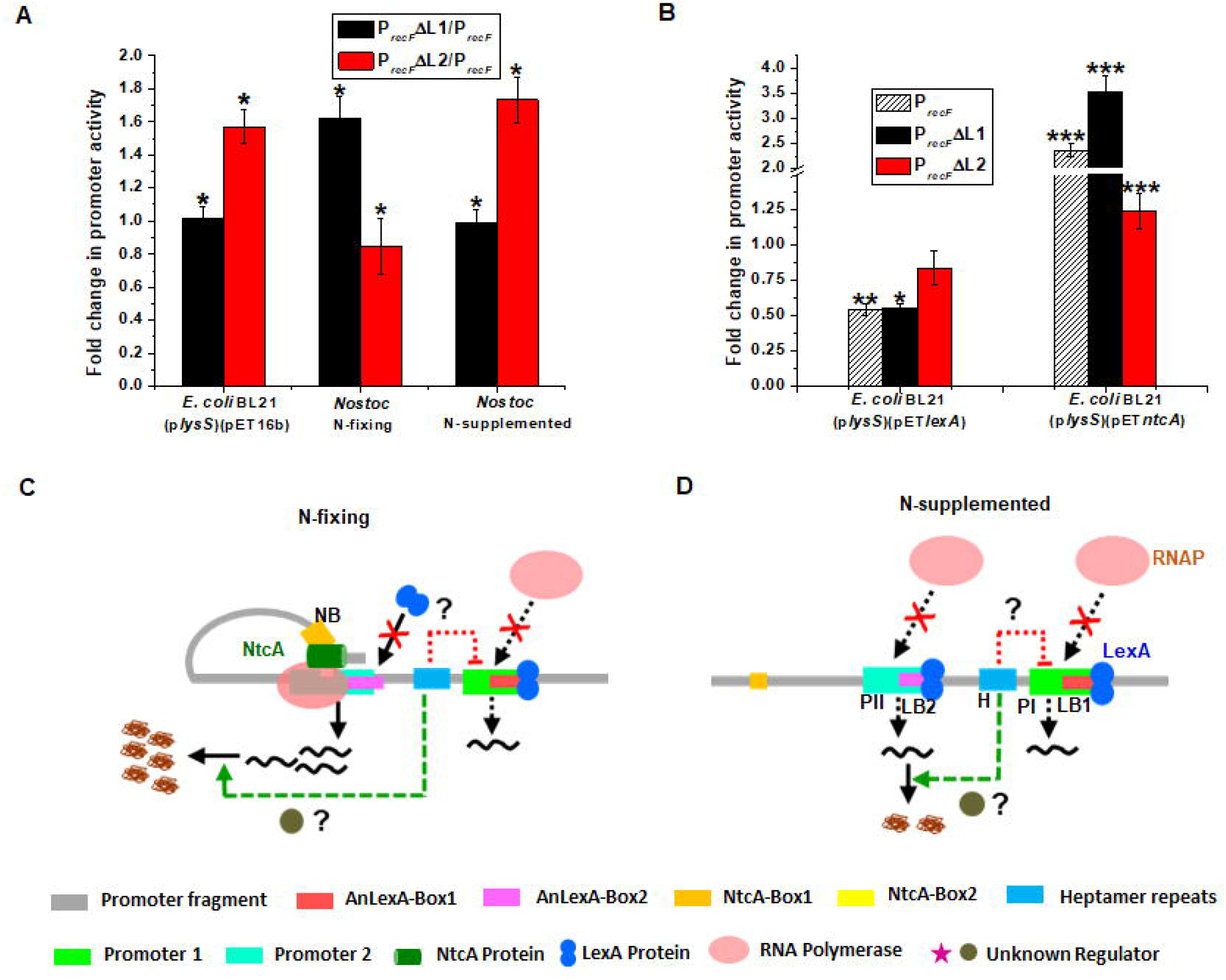
Regulation of *recF* gene of *Nostoc* 7120. (A and B) Fold change in promoter activity of (A) deletion constructs P_*recF*_ΔL1 and P_*recF*_ΔL2 with respect to full length P_*recF*_ in *E. coli* BL21(p*lysS*)(pET16b) cells in the absence of IPTG and in 3-day-old nitrogen-fixing and nitrogen-supplemented cultures of *Nostoc* 7120 as indicated below the bar graphs. (B) P_*recF*,_ P_*recF*_ΔL1 and P_*recF*_ΔL2 in *E. coli* cells overexpressing either LexA or NtcA as indicated below the bar graphs with respect to that in corresponding un-induced cells. LexA expression was induced with 1 mM IPTG for 2 h at 37 °C and NtcA with 1 mM IPTG for 16 h at 20 °C in *E. coli* BL21 cells. The absolute values of the activity of the full length promoters are as given in Fig. 1B. The average and s.d. values in correspond to experiments carried out as two biological and three technical replicates, and the * indicates p value (determined by t-test) as stated in legend to Fig. 1A. (C and D) Schematic representation of regulation of *recF* gene by different *cis*-elements and regulatory proteins under (C) N-fixing and (D) N-supplemented conditions in *Nostoc* 7120. The colour code of the different elements and proteins is shown below the figure. The dotted lines/arrows indicate partial transcription (black), repression (red) or activation (green).

In the presence of LexA (2 h after induction), the promoter activity was found to be repressed in the absence of either of the AnLexA-boxes in P_*recF*_, the fold-reduction being ∼1.8-fold for both P_*recF*_ and P_*recF*_ΔL1, and no change for P_*recF*_ΔL2) (Fig. 3A). Deletion of both the AnLexA boxes resulted in complete loss of promoter activity (data not shown) as it impaired both PI and PII promoters (Fig. 1C). Thus, it can be said that under *in vivo* conditions the down-regulation of promoter activity is through binding to AnLexA-Box2. Based on this, one would expect that the promoter activity of P_*recF*_ΔL2 would be higher than that of P_*recF*_ and P_*recF*_ΔL1 in *Nostoc* as well. However, this was true only in N-sup cultures wherein it increased by 1.75-fold, while it was slightly lower than that of P_*recF*_ under N-fix conditions (Fig. 3A), indicating involvement of NtcA in regulating the expression. Overexpression of *Nostoc* NtcA resulted in enhanced activity P_*recF*_ and P_*recF*_ΔL1 by 2.36-fold and 3.53-fold respectively, but no change for P_*recF*_ΔL2 in *E. coli* (Fig. 3B), which does not have the NtcA-Box (Fig. 1C). Thus, the positive contribution of NtcA in transcriptional up-regulation is lost under N-fix conditions in P_*recF*_ΔL2. While under N-sup conditions, wherein NtcA is not present, the increased activity is due to the de-repression of transcription due to the non-availability of AnLexA-Box2 for the binding of LexA. In fact, the absolute activity of P_*recF*_ΔL2 was similar under N-fix and N-sup conditions, thus confirming that the positive regulatory effect of NtcA is negated in the absence of the NtcA-Box in P_*recF*_ΔL2. The higher activity of P_*recF*_ΔL1 (which has NtcA-Box, AnLexA-Box2 and PII promoter) compared to P_*recF*_ under N-fix conditions, but near equal activity under N-sup conditions (Fig. 3A) indicates that (i) the positive regulatory effect of NtcA through PII negates the transcriptional repression through the binding of LexA to AnLexA-Box2, and (ii) confirms that PII is the preferred promoter, conforming to the results obtained in *E. coli*.

The third probable regulatory element detected upstream of the *recF* gene are the contiguous heptamer repeats (Figs. 1C, S1A). The *recF* gene in *Nostoc* is transcribed in the left direction, and hence the complementary sequence is shown in Fig. S1A, wherein the heptamer repeat is indicted as (TTCAAAA)_3_. However, in the genome it will be reflected as (TTTTGAA)_3_ upstream of the *recF*. A genome wide search revealed the presence of this heptamer as 3-4 contiguous repeats upstream (either within 100 bases of either the start codon or the predicted -10 promoter region) of 14 genes transcribed in the left orientation and 7 in right orientation (Table 1A), indicating it could have wider implications in gene regulation, at transcriptional, post-transcriptional or translational level. Though the *recF* transcript levels were low, RecF protein levels were found to be high in *Nostoc* 7120 (35). This is despite having a non-canonical S.D. sequence (ATCC) 12 bases upstream of ATG (Fig. S1) higher than the preferred distance of 4-8 bases. This raises the possibility of the involvement of the heptamer repeats in regulating translation. The role of heptamer repeats in regulating expression of downstream genes has not been very well worked out in bacteria. However, there are two reports indicating their possible role in gene regulation. The very first report was on a suggested role for the heptamer repeats in regulating transcription/translation binding slippage during reads in the *Beggiatoacaea* genome (47). In *Streptomyces natalensis*, the heptamer repeats have been shown to be important for DNA binding and regulating expression of *luxA* (48).

**Table 1A:** List of genes in *Nostoc* PCC7120 with a minimum of three contiguous repeats of the heptamer ‘TTTTGAA’ (Hept_L_) or ‘TTCAAAA’ (Hept_R_) upstream to the start codon.

| S.<br>No. | allX-N<sub>L</sub>-(TTTTGAA)<sub>n</sub> |  |  | S.<br>No. | (TTCAAAA)<sub>n</sub>-N<sub>R</sub>-alrY |  |  |
| --- | --- | --- | --- | --- | --- | --- | --- |
|  | n | allX | N<sub>L</sub> |  | n | alrY | N<sub>R</sub> |
| 1 | 3 | *all0341*(unknown) | 17 | 1 | 3 | *alr0608* (*nrtA*) | 92 |
| 2 | 4 | *all0420* (hypothetical) | 53 | 2 | 4 | *alr2054* (Aldo/ketoreductase) | 164 |
| 3 | 3 | *all0601*(hypothetical) | 183 | 3 | 3 | *alr2064* (hypothetical) | 172 |
| 4 | 4 | *all0769* (acetyl CoA synthetase) | 252 | 4 | 4 | *alr2119* (FeII transporter) | 93 |
| 5 | 3 | *all0980* (DDL)) | 251 | 5 | 4 | *alr2836* (glycosyltransferase) | 152 |
| 6 | 3 | *all0988* (NagA) | 86 | 6 | 3 | *alr2967* (unknown) | 89 |
| 7 | 3 | *all1089* (hypothetical) | 114 | 7 | 4 | *alr4660* (hypothetical) | 109 |
| 8 | 4 | *all1091* (unknown) | 02 |  |  |  |  |
| 9 | 3 | *all1291* (*cynS*) | 136 |  |  |  |  |
| 10 | 4 | *all2020* (ABC transporter ATP-binding protein) | 165 |  |  |  |  |
| 11 | 4 | *all2220* (hypothetical) | 61 |  |  |  |  |
| 12 | 4 | *all2870* (unknown) | 301 |  |  |  |  |
| 13 | 3 | *all3374* (*recF*) | 61 |  |  |  |  |
| 14 | 3 | *all5057* (*accB*) | 102 |  |  |  |  |

allX refers to genes in left orientation and alrY to genes in right orientation. ‘n’ refers to number of heptamer repeats, N<sub>L</sub> and N<sub>R</sub> is the distance between the left/right gene and start of heptamer, shown schematically below.

In a nutshell, the transcription of *recF* is initiated from PII promoter, which is (i) activated through the binding of NtcA to NtcA-Box and (ii) repressed through the binding of LexA to either of the two AnLexA-boxes, extent of repression being higher upon binding to AnLexA-Box2 (Figs. 3C, D). As shown in Fig. 3C, under N-fix conditions, the binding of the NtcA enhances the transcription from PII and prevents the binding of LexA to AnLexA-Box2, allowing the repression through LexA to occur only through binding to AnLexA-Box1, which functions as a weaker repressor. On the other hand, under N-sup conditions as shown in Fig. 3D, LexA can bind to both the AnLexA-boxes due to absence of NtcA. Since, the AnLexA-Box1 overlaps with PII, it significantly represses the transcription (Fig. 3D), resulting in lower transcript levels of *recF* under N-sup compared to N-fix conditions (Fig. 1B). Though the exact role of the heptamer repeats could not be delineated, we speculate that it could be involved in either transcriptional/ post-transcriptional/ translational regulation or both of the *recF* gene. At transcriptional level, the heptamer repeats could be interfering with the binding of RNA Polymerase to PI (Figs. 3C, D), thus accounting for the preferential use of PII in spite of it being weaker than PI (Fig. 3A, B). In *Nostoc* 7120, though the transcript levels of *recF* were low, the protein levels were comparatively higher (35), despite the presence of a distant and non-canonical S.D. sequence in case of *recF* (Fig. S1A). Thus, it has also been speculated that the heptamer repeats upstream of the *recF* gene may be involved in enhancing the translation (Figs. 3C, D), as has also been predicted for the heptamer repeats of *Beggiatoacaea* genome (47).

### 3.4 Regulation of expression of *recO* gene

The *recO* gene of *Nostoc* 7120 is flanked by two genes transcribed in opposite directions, and hence is regulated by its own promoter unlike the autoregulation by RNaseIII promoter observed in other bacteria. The region upstream of the *recO* gene of *Nostoc* 7120 is characterised by a single promoter which overlaps with AnLexA-Box2 and NtcA-Box, and a heptamer repeat ‘GAC(C/T)AAT repeated 8 times contiguously downstream to AnLexA-Box1 and overlapping with the first two bases of the start codon (Fig. 1C, Fig. S1B). In case of P_*recO*_, the promoter activity was found to be unaffected by deletion of either the heptamer repeats alone (ΔH) or heptamer with AnLexA-Box1 (ΔHL1) or upon mutation of AnLexA-Box2 (mutL2) (Fig. 4A). However, in the presence of *Nostoc* LexA, a 2.22-fold and 1.78-fold decrease in promoter activity was observed for P_*recO*_ΔH and P_*recO*_ΔHL1 respectively as compared to no change for P_*recO*_ and P_*recO*_mutL2 (Fig. 4B). Higher repression by LexA observed for P_*recO*_ΔH compared to P_*recO*_ΔHL1 in *E. coli* (Fig. 4B) indicates that both the AnLexA-boxes are functional and have an additive effect in repressing the promoter activity. Similarly, the deletion of the heptamer repeats resulted in increased promoter activity in the presence of NtcA, the fold-increase being 2.51 and 2.87 for P_*reco*_ΔH and P_*recO*_ΔHL1 respectively, compared to 1.38-fold for P_*recO*_ (Fig. 4B). Thus, the repression by LexA and the activation by NtcA of the *recO* promoter were achieved only in the absence of the heptamer repeats in *E. coli*, suggesting a regulatory role for the heptamer repeats as well.

**Figure 4.**
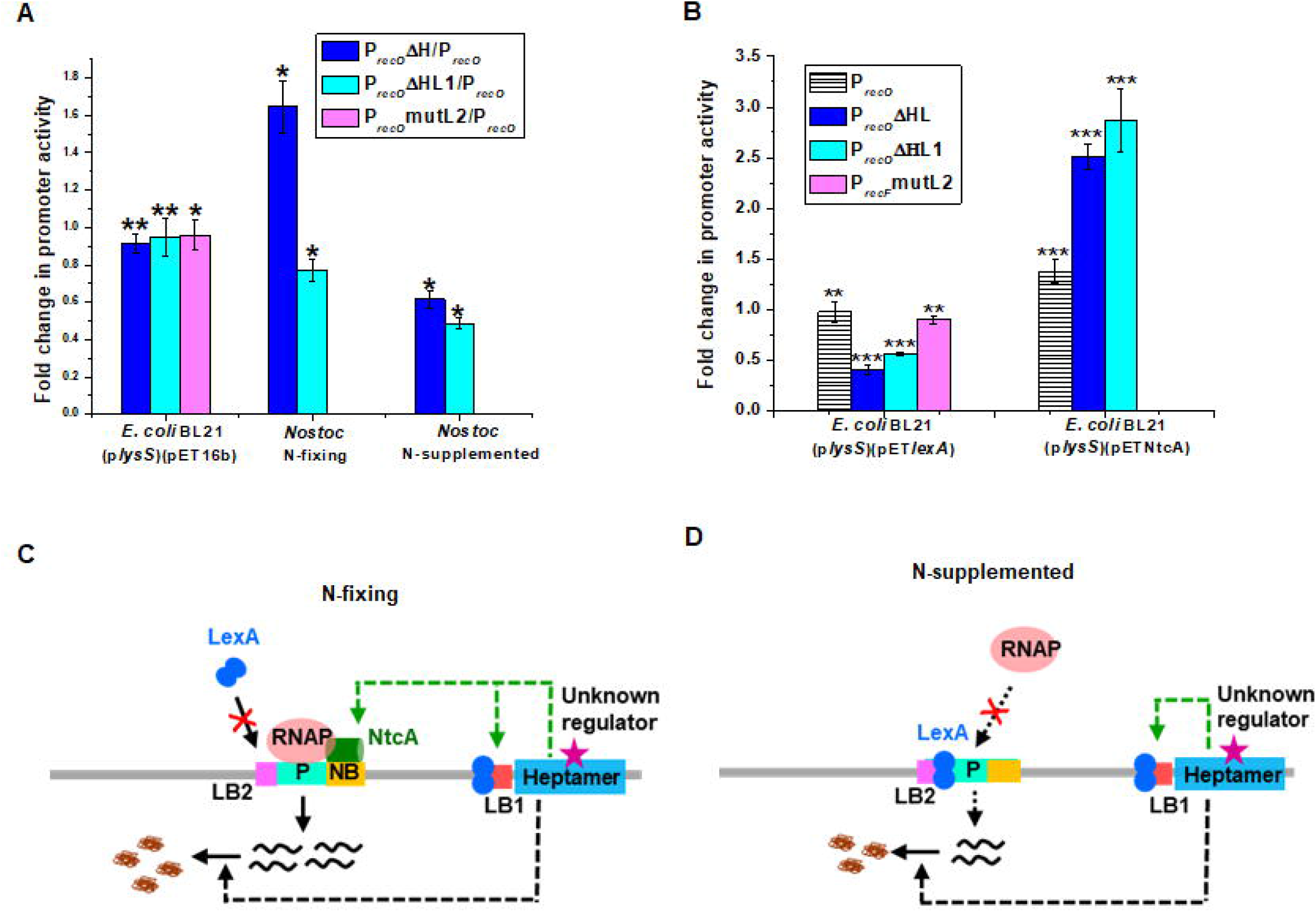
Regulation of *recO* gene of *Nostoc* 7120. (A and B) Fold change in promoter activity of (A) deletion constructs P_*recO*_ΔH, P_*recO*_ΔHL1 and P_*recO*_mutlL2 with respect to full length P_*recO*_ in *E. coli* BL21(p*lysS*)(pET16b) cells in the absence of IPTG and in 3-day-old N-fix and N-sup cultures of *Nostoc* 7120 as indicated below the bar graphs. (B) P_*recO*,_ P_*recO*_ΔH and P_*recO*_ΔHL1 in *E. coli* cells in the presence of either LexA or NtcA compared to that in un-induced cells overexpressing either LexA or NtcA as indicated below the bar graphs with respect to that in corresponding un-induced cells. Other details are as given in legend for Fig. 3A, B.. (C and D) Schematic representation of regulation of *recO* gene in *Nostoc* 7120 by different *cis*-elements and regulatory proteins under (C) N-fix and (D) N-sup conditions. Other details are as indicated in legend to Fig. 3C, D.

Even in *Nostoc* 7120, deletion of the heptamer repeats resulted in the modulation of promoter activity. Compared to P_*recO*_, the activity of P_*recO*_ΔH increased 1.6-fold under N-fix, but decreased 1.5-fold under N-sup conditions, while that of P_*recO*_ΔHL1 decreased under both the conditions, the fold decrease being much higher under N-sup conditions (2.06-fold) compared to that under N-fix conditions (1.29-fold) (Fig. 4A). The up-regulation of promoter activity of full length (P_*recO*_) as well as deletion constructs (P_*recO*_ΔH and P_*recO*_ΔHL1) in the presence of NtcA in *E. coli* (Fig. 4B) concurs with the higher promoter activity of all three constructs under N-fix conditions compared to N-sup conditions in *Nostoc* (Figs. 1B, 4A). Also, the positive regulatory effect of NtcA was higher in the absence of the heptamer repeats, indicating a regulatory role for the heptamer repeats in the binding of NtcA to the NtcA-Box GTA-N_10_-C overlapping with the promoter region of *recO* (Fig. S1B).

It can be speculated that the secondary structure arising out of the heptamer repeats may prevent the access for binding of LexA under *in vivo* conditions in *E. coli*. Since, in *Nostoc* 7120, the transcriptional repression of *recO* by LexA is observed (Fig. 1A), it can be further speculated that a *Nostoc* specific protein interacting with the heptamer sequence may be involved in gaining accessibility to the LexA protein to cause repression through binding to the AnLexA-boxes. The heptamer repeat was specific not only for the *recO* gene, but was found upstream (within 30 bases of the start codon) of at least 13 genes in *Nostoc* 7120 (Table 1B), and thus could be a more general gene regulatory element.

**Table 1B.**
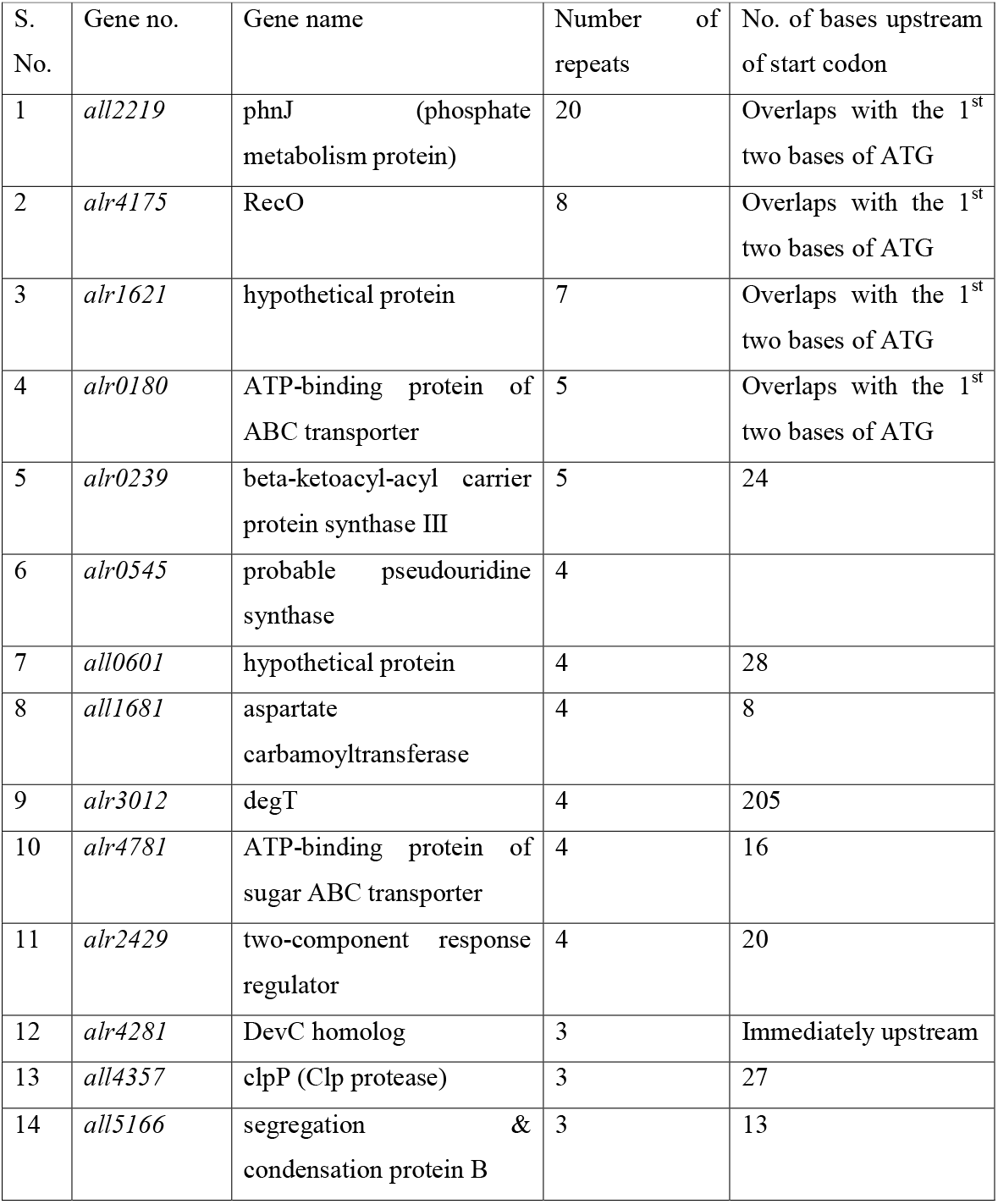
List of genes with a minimum of three contiguous repeats of the heptamer GA(C/T)AAT.

| S. No. | Gene no. | Gene name | Number of repeats | No. of bases upstream of start codon |
| --- | --- | --- | --- | --- |
| 1 | *all2219* | phnJ (phosphate metabolism protein) | 20 | Overlaps with the 1<sup>st</sup> two bases of ATG |
| 2 | *alr4175* | RecO | 8 | Overlaps with the 1<sup>st</sup> two bases of ATG |
| 3 | *alr1621* | hypothetical protein | 7 | Overlaps with the 1<sup>st</sup> two bases of ATG |
| 4 | *alr0180* | ATP-binding protein of ABC transporter | 5 | Overlaps with the 1<sup>st</sup> two bases of ATG |
| 5 | *alr0239* | beta-ketoacyl-acyl carrier protein synthase III | 5 | 24 |
| 6 | *alr0545* | probable pseudouridine synthase | 4 |  |
| 7 | *all0601* | hypothetical protein | 4 | 28 |
| 8 | *all1681* | aspartate carbamoyltransferase | 4 | 8 |
| 9 | *alr3012* | degT | 4 | 205 |
| 10 | *alr4781* | ATP-binding protein of sugar ABC transporter | 4 | 16 |
| 11 | *alr2429* | two-component response regulator | 4 | 20 |
| 12 | *alr4281* | DevC homolog | 3 | Immediately upstream |
| 13 | *all4357* | clpP (Clp protease) | 3 | 27 |
| 14 | *all5166* | segregation & condensation protein B | 3 | 13 |

As shown in Fig. 4C, the binding of NtcA to NtcA-Box under N-fix conditions probably precludes the binding of LexA to AnLexA-Box2, thereby allowing higher access to RNAP to bind to the promoter resulting in higher expression of *recO* under nitrogen-fixing conditions. While under N-sup conditions, the binding of LexA to AnLexA-box2 decreases the accessibility of promoter for the binding of RNAP, leading to lower transcription (Fig. 4D). It is speculated that a protein may be binding the heptamer repeats in *Nostoc* 7120, thereby allowing positive regulation by NtcA and negative regulation by LexA of the *recO* promoter (Figs. 4C, D). Thus, it is further speculated that the cellular ratio of NtcA and LexA along with the yet to be identified interacting partner of the heptamer repeats would be dictating the expression of *recO* under different growth conditions. Being in close vicinity of the start codon, in fact overlapping with it in 4 cases, it may also play a role in post-transcriptional control. This could account for the low levels of RecO protein in *Nostoc* 7120 (35). Hence, the identification of the protein binding to the heptamer repeat could hold the key for the intricate regulation of expression of *recO*, and also the other genes which have these heptamer repeats in close vicinity of their promoter or start codon (Table 1B).

### 3.5 Regulation of expression of *recR* gene

Both the promoters upstream of *recR* (Figs. 1C, S1C) seem to be contributing towards its expression in the heterologous host *E. coli*, as the basal promoter activity decreased 1.5-fold in the absence of either of the promoters (Fig. 5A) which are disrupted upon deletion of the closely placed corresponding AnLexA-Box (Fig. 1C). In the presence of LexA (2 h after induction), the promoter activity was found to be repressed in the absence of either of the AnLexA-boxes in P_*recR*_, with the fold-reduction being 3.2-fold and 3.5-fold respectively for P_*recR*_ and P_*recR*_ΔL1, and no change for P_*recR*_ΔL2 (Fig. 5A). No activity was observed when both the AnLexA boxes were deleted from P_*recR*_ (data not shown), which indicated that it has only two functional promoters PI and PII.

**Figure 5.**
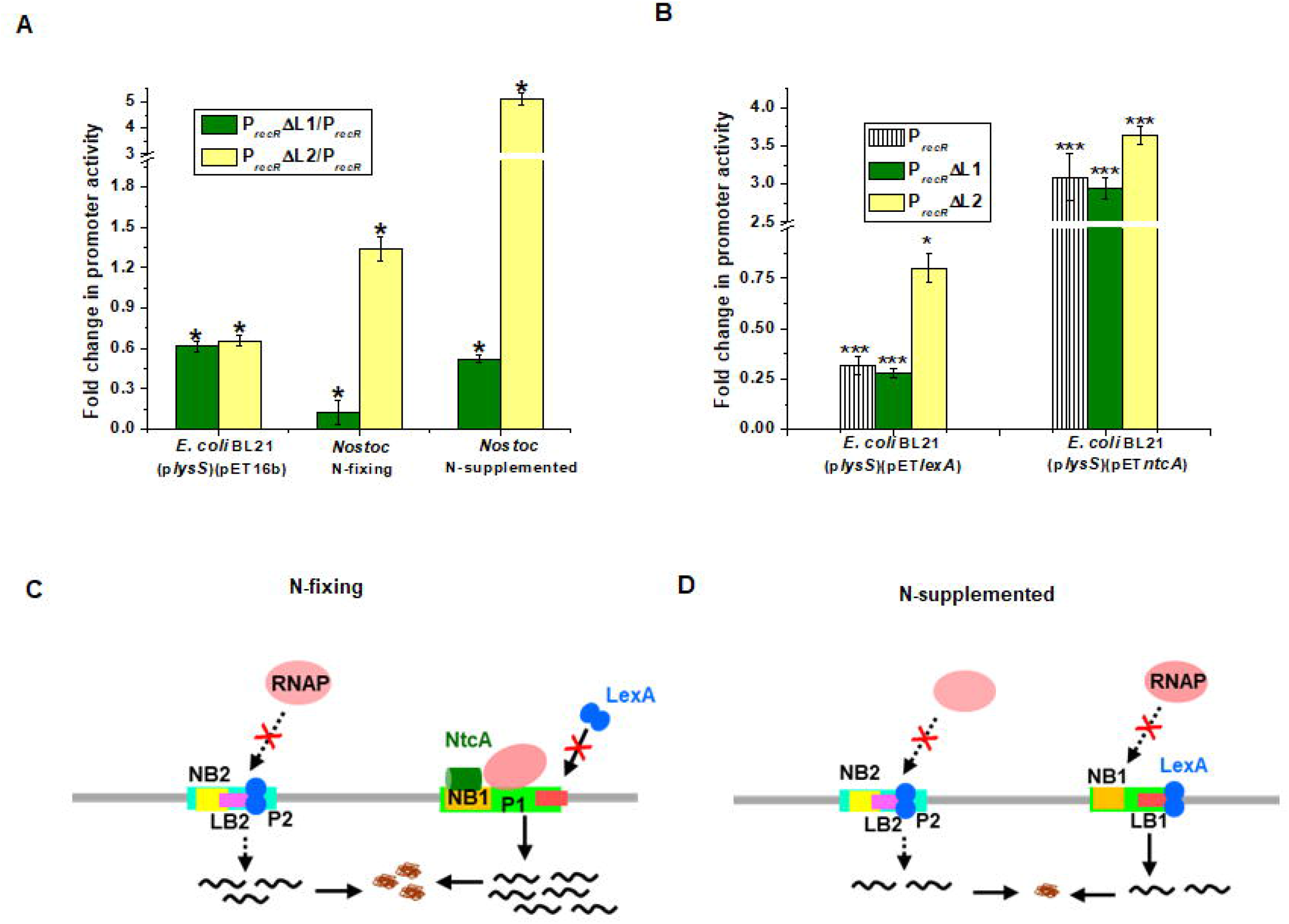
Regulation of *recR* gene of *Nostoc* 7120. (A and B) Fold change in promoter activity of (A) deletion constructs P_*recR*_ΔL1 and P_*recR*_ΔL2 with respect to P_*recR*_ in *E. coli* BL21(p*lysS*)(pET16b) cells in the absence of IPTG and in 3-day-old N-fix and N-sup cultures of *Nostoc* 7120 as indicated below the bar graphs. (B) P_*recR*,_ P_*recR*_ΔL1 and P_*recR*_ΔL2 in *E. coli* cells in the presence of either LexA or NtcA compared to that in un-induced cells overexpressing either LexA or NtcA as indicated below the bar graphs with respect to that in corresponding un-induced cells. Other details are as given in legend for Fig. 3A, B.. (C and D) Schematic representation of regulation of *recR* gene in *Nostoc* 7120 by different *cis*-elements and regulatory proteins under (C) N-fix and (D) N-sup conditions. Other details are as indicated in legend to Fig. 3C, D.

P_*recR*_ΔL1 exhibited 7.9-fold and 2-fold lower activity under N-fix and N-sup conditions respectively (Fig. 5A). On the other hand, P_*recR*_ΔL2 exhibited 1.34-fold and 7.3-fold higher activity under N-fix and N-sup conditions respectively compared to P_*recR*_ (Fig. 5A). This is despite the fact that both the NtcA-boxes are intact in P_*recR*_ΔL1 (Fig. 1C). In the presence of NtcA in *E. coli*, the increase in activity was ∼3-fold for P_*recR*_ and P_*recR*_ΔL1 and ∼3.64-fold for P_*recR*_ΔL2 (Fig. 5B). This clearly indicates that the up-regulation of *recR* transcription is through NtcA-Box1 and PI. While both the NtcA-boxes had the sequence GTA-N_10_-C, the base preceding ‘C’ was ‘A’ in NtcA-Box1 and ‘G’ in NtcA-Box2, and based on earlier report that the occurrence of ‘T’ and ‘A’ preceding ‘C’ in NtcA-box is higher than that of ‘G’ (31), it can be stated that *recR* has only one functional NtcA-Box corresponding to NtcA-Box1.

Since, only NtcA-Box1 (NB1) is active in *recR*, the binding of LexA dimer to AnLexA-Box2 (LB2) results in decreased transcription from PII promoter irrespective of the N-status (Figs. 5C, D). Under N-fix condition, the binding of NtcA to NB1 precludes the binding of LexA to LB1 and thus there is enhanced transcription from PI (Fig. 5C). Under N-sup conditions, where there is no NtcA, transcription through PI is also inhibited (Fig. 5D) leading to lower transcription and promoter activity under N-sup conditions compared to N-fix conditions (Fig. 1A). The expression of *recR* is also regulated at the translational level as seen by extremely low basal levels of this protein in *Nostoc* 7120 (35), through a rare initiation codon ‘GTT’. Analysis of the 2° structure of the tRNA-fMet (position 5019099..5019172 on *Nostoc* 7120 genome) revealed the presence of the anticodon corresponding to the ‘GTT’ codon at the end of the anticodon loop (Fig. S3), and thus would be able to bind and initiate transcription from ‘GTT’ taking advantage of a strong SD sequence (Fig. S1C), though at a relatively lower rate compared to that for common initiation codons such as ‘ATG’ and ‘GTG’.

### 3.6 Regulation of expression of *recF, recO* and *recR* genes during post-irradiation recovery (PIR) in *Nostoc*

In order to assess the physiological relevance of the *recF, O, R* genes especially with respect to DA repair, their expression (transcript and promoter activity) was monitored upon exposure to γ-irradiation and during post-irradiation recovery (PIR). Under N-fix conditions, the activity of P_*recF*_ decreased till 22 h of PIR followed by a 1.5 and 2-fold increase at the end of 32 h and 45 h of PIR respectively (Fig. 6A). Under N-sup conditions, activity of P_*recF*_ showed a 3-fold decrease immediately after γ-radiation, and thereafter showed a slow increase reaching activity equal to sham-irradiated controls at the end of 45 h (Fig. 6A). Activity of P_*recO*_ decreased 1.8-fold under nitrogen-fixing conditions and increased 1.4-fold under nitrogen-supplemented conditions immediately after γ-radiation (Fig. 6B). Thereafter during PIR, the activity of P_*recO*_ was similar to that of sham-irradiated control (SIC) irrespective of the N-status (Fig. 6B). Activity of P_*recR*_ decreased 3.6-fold immediately after γ-radiation under N-sup conditions, while during PIR, the values were similar to that of SIC irrespective of N-status (Fig. 6C).

**Figure 6.**
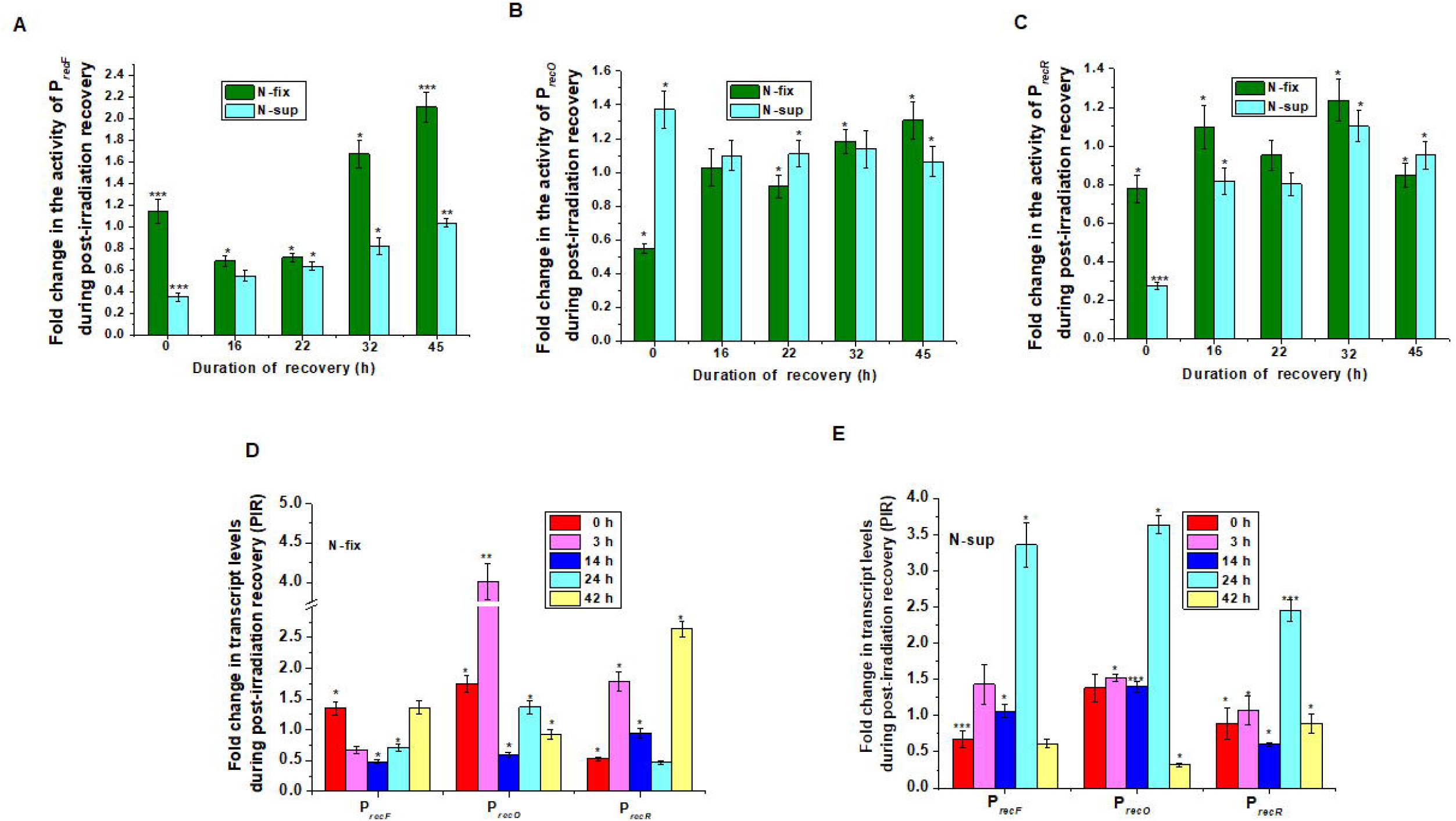
Expression of *recF, recO* and *recR* during post-irradiation recovery. Three-day-old of *Nostoc* PCC7120 exposed to 6 kGy γ-irradiation (or corresponding sham-irradiation) were allowed to recover under normal growth conditions. (A-C) Fold change in promoter activity during PIR compared to that of sham-irradiated controls (SIC) is shown for (A) P_*recF*_, (B) P_*recO*_ and (C) P_*recR*_ under nitrogen-fixing (N-fix) and nitrogen-supplemented (N-sup) conditions. (D and E) Transcription profile using qRT-PCR analysis was carried for *recF, recO* and *recR* genes from RNA isolated at the end of 0, 3, 14, 24 and 42 h of the recovery period from (D) nitrogen-fixing and (E) nitrogen-supplemented cultures and expressed as fold-change during PIR with respect to their corresponding sham-irradiated (SIC) recovered cultures. Other details are as described in legend to Fig. 1A.

Transcript expression profile of *recF, recO* and *recR* genes was monitored at different time points of PIR up to 42 h by qRT-PCR, and compared with the corresponding SIC. Under nitrogen-fixing conditions, *recF* transcript levels increased 1.36-fold immediately after γ-irradiation, followed by decreased expression by ∼2-fold at 14 h PIR and thereafter increasing to levels comparable to that immediately after irradiation at the end of 42 h of PIR (Fig. 6D). Under N-sup conditions, *recF* transcript levels decreased by 1.47-fold immediately after γ-irradiation, and thereafter increasing, reaching maximal fold induction (3.3-fold) at the end of 24 h of PIR (Fig. 6E). The *recO* transcript levels upon γ-irradiation and during PIR were found to be similar or higher than in SIC, with the exception of 2-fold lower transcript levels at 14 h PIR under N-fix conditions (Fig. 6D) and 3.3-fold lower transcript levels at 42 h of PIR under N-sup conditions (Fig. 6E). Maximal induction of *recO* transcripts during PIR was observed at 3 h (3.2-fold) and 24 h (3.5-fold) under N-fix and N-sup conditions respectively (Figs. 6D, E). Transcript levels of *recR* immediately after γ-irradiation decreased only under N-fix conditions (∼2-fold) (Figs. 6D, E). During PIR under N-fix conditions, it was found to increase by 2-fold at 3 h PIR, followed by a steady decrease and then an increase by 2.5-fold at 42 h PIR (Fig. 6D). Under N-sup conditions, the decrease was by 2-fold at 14 h of PIR and exhibited maximal increase by 2.5-fold at 24 h of PIR (Fig. 6E). Both NtcA and LexA proteins have been shown to be repressed upon exposure to radiation in cyanobacteria, that of NtcA in *Arthrospira* (3) and LexA in *Nostoc* (27). The recovery of the synthesis and activity of the two proteins during post-irradiation recovery would regulate the expression of the *recF, recO* and *recR* genes in *Nostoc* 7120 and thus account for the differences observed in their expression profile under nitrogen-fixing and nitrogen-supplemented conditions (Fig. 6). The heptamer repeats and its co-activator/repressor would also contribute to the regulation of expression of *recF* and *recO* genes during PIR.

## 4. CONCLUSION

Expression of *recF, recO* and *recR* genes in *Nostoc* 7120 is regulated at different levels through interplay of regulatory proteins and *cis-*elements in the promoter. Involvement of the multiple transcriptional regulatory elements and proteins, *viz* negative regulator LexA, positive regulator NtcA, heptamer repeats and translational regulatory elements, *viz* non-canonical S.D. sequence and translational start codon help fine tune the expression of these genes not only under normal growth conditions, but also in response to DNA damage inducing stresses such as γ-radiation, which is essential to ensure effective repair of DSBs and recovery of these cells post-radiation. Of these, the role of NtcA in regulating DNA repair proteins as part of its global regulatory role is being reported for the first time in cyanobacteria. Cyanobacterial genome is replete with several repeat sequences especially heptamer repeats. A thorough investigation on these in terms of their role in DNA replication, transcriptional and translational regulation will throw new insights into the complex gene regulatory networks existing in cyanobacteria.

## ACKNOWLEDGEMENT

Mitali Pradhan acknowledges DAE/HBNI for JRF fellowship. Sarita Pandey thanks DST (DST/INSPIRE/04/2016/000768), New Delhi for INSPIRE Faculty Award for funding. MP and SP thanks MBD, BARC for providing research facilities.

## FIGURE LEGENDS

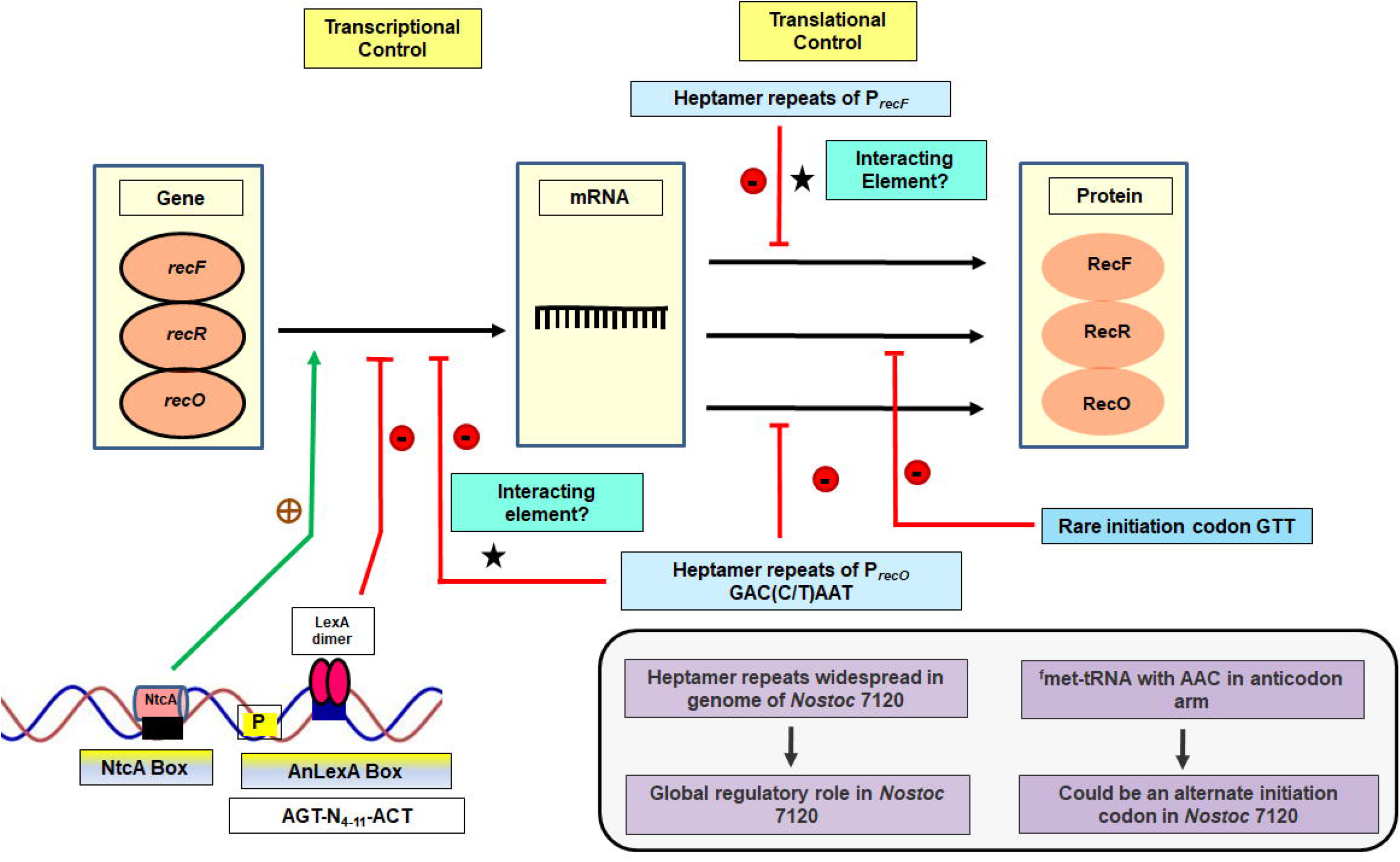

## REFERENCES

1. Brock, T.D., (1973). Lower pH limit for the existence of blue-green algae: evolutionary and ecological implications. Science 179, 480–483.

2. Billi, D., Friedmann, E.I., Hofer, K.G., Caiola, M.G., Ocampo-Friedmann, R,. (2000). Ionizing-radiation resistance in the desiccation-tolerant cyanobacterium Chroococcidiopsis. Appl. Environ. Microbiol. 66, 1489–92.

3. Badri, H., Monsieurs, P., Coninx, I., Nauts, R., Wattiez, R., Leys, N., (2015) Temporal gene expression of the cyanobacterium Arthrospira in response to Gamma Rays. PLoS ONE 10, e0135565.

4. Singh, H., Anurag, K., Apte, S.K., (2013). High radiation and desiccation tolerance of nitrogen-fixing cultures of the cyanobacterium Anabaena sp. strain PCC 7120 emanates from genome/proteome repair capabilities. Photosyn. Res. 118, 71–81.

5. Rajaram, H., Kumar, A., Kirti, A., Pandey, S., (2020). Double strand break (DSB) repair in Cyanobacteria: Understanding the process in an ancient organism. DNA Repair 95, 102942.

6. Zahradka, K., Slade, D., Bailone, A., Sommer, S., Averbeck, D., Petranovic, M., Lindner, A. B., Radman, M., (2006). Reassembly of shattered chromosomes in Deinococcus radiodurans. Nature 443, 569–573.

7. Bentchikou, E., Servant, P., Coste, G., Sommer, S., (2010). A major role of the RecFOR pathway in DNA double-strand-break repair through ESDSA in Deinococcus radiodurans. PLoS Genetics 6, e1000774.

8. Kowalczykowski, S.C., Dixon, D.A., Eggleston, A.K., Lauder, S.D., Rehrauer, W.M., (1994). Biochemistry of homologous recombination in Escherichia coli. Microbiol. Rev. 58, 401–465.

9. Kuzminov, A., (1999). Recombinational repair of DNA damage in Escherichia coli and bacteriophage lambda. Microbiol. Mol. Biol. Rev. 63, 751–813.

10. Armengod, M.E., (1982). recF-dependent recombination as a SOS function. Biochimie 64, 629–632.

11. Walker, G.C., (1984). Mutagenesis and inducible responses to deoxyribonucleic acid damage in Escherichia coli. Microbiol. Rev. 48, 60–93.

12. Sandler, S.J., Clark, A.J., (1994). Mutational analysis of sequences in the recF gene of Escherichia coli K-12 that affect expression. J. Bacteriol. 176, 4011–4016.

13. Armengod, M.E., Lambíes, E. (1986). Overlapping arrangement of the recF and dnaN operons of Escherichia coli; positive and negative control sequences. Gene 43, 183–196.

14. Pérez-Roger, I., García-Sogo, M., Navarro-Aviñó, J.P., López-Acedo, C., Macián, F., Armengod, M.E., (1991). Positive and negative regulatory elements in the dnaA-dnaN-recF operon of Escherichia coli. Biochimie 73, 329–334.

15. Villarroya, M., Pérez-Roger, I., Macián, F., & Armengod, M. E. (1998). Stationary phase induction of dnaN and recF, two genes of Escherichia coli involved in DNA replication and repair. EMBO J. 17(6), 1829–1837. https://doi.org/10.1093/emboj/17.6.1829

16. Badran, H., Venkatesh, T.V., Kunnimalaiyaan, M., Sharma, N., Das, H.K., (1995). Molecular characterization of the Azotobacter vinelandii recF gene. Gene 162, 47–51.

17. Anderson, P.E., Matsunaga, J., Simons, E.L., Simons, R.W., (1996). Structure and regulation of the Salmonella typhimurium rnc-era-recO operon. Biochimie 78, 1025–1034.

18. Matsunaga, J., Dyer, M., Simons, E.L., Simons, R.W., (1996). Expression and regulation of the rnc and pdxJ operons of Escherichia coli. Mol. Microbiol. 22, 977–989.

19. Zuber, M., Hoover, T.A., Powell, B.S., Court, D.L. (1994). Analysis of the rnc locus of Coxiella burnetii. Mol. Microbiol. 14, 291–300.

20. Powell, B., Peters, H.K., Nakamura, Y., Court, D., (1999). Cloning and analysis of the rnc-era-recO operon from Pseudomonas aeruginosa. J. Bacteriol. 181, 5111–5113.

21. Haraszthy, V.I., Jordan, S.F., Zambon, J.J., (2006). Identification of Fur-regulated genes in Actinobacillus actinomycetemcomitans. Microbiol (Reading) 152, 787–796.

22. Beaume, N., Pathak, R., Yadav, V.K., Kota, S., Misra, H S., Gautam, H.K., Chowdhury, S., (2013). Genome-wide study predicts promoter-G4 DNA motifs regulate selective functions in bacteria: radioresistance of D. radiodurans involves G4 DNA-mediated regulation. Nucleic Acids Res. 41, 76–89.

23. Flower, A.M., McHenry, C.S., (1991). Transcriptional organization of the Escherichia coli dnaX gene. J. Mol. Biol. 220, 649–658.

24. Yeung, T., Mullin, D.A., Chen, K.S., Craig, E.A., Bardwell, J.C., Walker, J.R., (1990). Sequence and expression of the Escherichia coli recR locus. J. Bacteriol. 172, 6042–6047.

25. Herrero, A., Muro-Pastor, A.M,, Flores, E. (2001) Nitrogen control in cyanobacteria. J. Bacteriol. 183, 411–425.

26. Li, S., Xu, M., Su, Z., (2010) Computational analysis of LexA regulons in Cyanobacteria. BMC Genomics 11, 527.

27. Kumar, A., Kirti, A., Rajaram, H., (2018). Regulation of multiple abiotic stress tolerance by LexA in the cyanobacterium Anabaena sp. strain PCC7120. Biochim. Biophys. Acta. Gene Reg. Mech. 1861, 864–877.

28. Espinosa, J., Rodríguez-Mateos, F., Salinas, P., Lanza, V.F., Dixon, R., de la Cruz, F., Contreras, A., (2014). PipX, the coactivator of NtcA, is a global regulator in cyanobacteria. Proc. Natl. Acad. Sci. USA 111, E2423–E2430.

29. Mitschke, J., Vioque, A., Haas, F., Hess, W.R., Muro-Pastor, A.M., (2011) Dynamics of transcriptional start site selection during nitrogen stress-induced cell differentiation in Anabaena sp. PCC7120. Proc Natl Acad Sci U S A. 108, 20130–20135.

30. Giner-Lamia, J., Robles-Rengel, R., Hernández-Prieto, M.A., Muro-Pastor, M.I., Florencio, F.J., Futschik, M.E., (2017). Identification of the direct regulon of NtcA during early acclimation to nitrogen starvation in the cyanobacterium Synechocystis sp. PCC 6803. Nucleic Acids Res. 45, 11800–11820.

31. Picossi, S., Flores, E., Herrero, A., (2014) ChIP analysis unravels an exceptionally wide distribution of DNA binding sites for the NtcA transcription factor in a heterocyst-forming cyanobacterium. BMC Genomics 15, 22.

32. Srivastava, A., Kumar, A., Biswas, S., Srivastava, V., Rajaram, H., Mishra, Y., (2022) Regulatory role of LexA in modulating photosynthetic redox poise and cadmium stress tolerance in the cyanobacterium, Anabaena sp. PCC7120. Environ. Expt. Bot. 195, 104790.

33. Domain, F., Houot, L., Chauvat, F., Cassier-Chauvat, C., (2004). Function and regulation of the cyanobacterial genes lexA, recA and ruvB: LexA is critical to the survival of cells facing inorganic carbon starvation. Mol Microbiol. 53, 65–80.

34. Kizawa, A., Kawahara, A., Takimura, Y., Nishiyama, Y., Hihara, Y., (2016). RNA-seq Profiling Reveals Novel Target Genes of LexA in the cyanobacterium Synechocystis sp. PCC 6803. Front Microbiol. 7, 193.

35. Pandey, S., Kumar, A., Kirti, A., Gupta, G.D., Rajaram, H., (2021). Rec(F/O/R) proteins of the nitrogen-fixing cyanobacterium Nostoc PCC7120: In silico and expression analysis. Gene 788, 145663.

36. Castenholz, RW., (1988) Culturing methods for cyanobacteria. Methods Enzymol.167, 68–93.

37. Kirti, A., Rajaram, H., Apte, S.K., (2013). Characterization of two naturally truncated, Ssb-like proteins from the nitrogen-fixing cyanobacterium, Anabaena sp. PCC7120. Photosyn. Res. 118, 147–154.

38. Mackinney, G., (1941). Absorption of light by chlorophyll solutions. J. Biol. Chem. 140, 315–322.

39. Yoon, H.S., Golden, J.W., (1998). Heterocyst pattern formation controlled by a diffusible peptide. Science 282, 935–938.

40. Raghavan, P.S., Rajaram, H., Apte, S.K. (2011) Nitrogen status dependent oxidative stress tolerance conferred by overexpression of MnSOD and FeSOD proteins in Anabaena sp. strain PCC7120. Plant. Mol. Biol. 77, 407–417.

41. Kirti, A., Kumar, A., Rajaram, H., (2017). Differential regulation of ssb genes in the nitrogen-fixing cyanobacterium, Anabaena sp. strain PCC7120. J. Phycol. 53, 322–332.

42. Livak, K.J., Schmittgen, T.D., (2001). Analysis of relative gene expression data using real-time quantitative PCR and the 2− ΔΔCT method. Methods 25, 402–408.

43. Schneider, C.A., Rasband, W.S., Eliceiri, K.W. (2012). NIH Image to ImageJ: 25 years of image analysis. Nature Methods 9, 671–675.

44. Rajaram, H., Apte, S.K., (2010). Differential regulation of groESL operon expression in response to heat and light in Anabaena. Arch. Microbiol. 192, 729–738.

45. Solovyev V., Salamov, A., (2011). Automatic annotation of microbial genomes and metagenomic sequences. Metagenomics and Its Applications in Agri., Biomed. Environ. Studies; Li, RW, Ed, 61–78.

46. Wei, Y., Xia, X., (2019). Unique Shine-Dalgarno Sequences in Cyanobacteria and Chloroplasts Reveal Evolutionary Differences in Their Translation Initiation. Genome Biol. Evol. 11, 3194–3206.

47. MacGregor B.J., (2015) Abundant intergenic TAACTGA direct repeats and putative alternate RNA polymerase β ′ subunits in marine beggiatoaceae genomes: possible regulatory roles and origins. Front. Microbiol. 6, 1397.

48. Barreales, E.G., Vicente, C.M., de Pedro, A., Santos-Aberturas, J., Aparicio, J.F., (2018). Promoter engineering reveals the importance of heptameric direct repeats for DNA binding by Streptomyces antibiotic regulatory protein - large ATP-binding regulator of the LuxR family (SARP-LAL) regulators in Streptomyces natalensis. Appl. Environ. Microbiol. 84, e00246.

