## Supplementary Tables for "NtcA, LexA and heptamer repeats involved in the multifaceted regulation of DNA repair genes *recF, recO* and *recR* in the cyanobacterium *Nostoc* PCC7120"

**Supplementary Table S1A: List of primer sequences used for cloning**

| **Sr No** | **Primer Name** | **Primer Sequence*** |
| --- | --- | --- |
| 1 | P*_recF_*Fwd | GACGAGCTCTCCTGTCTCAAATCCTTATGT |
| 2 | P*_recF_*Rev | GACGGTACCAAAAAACAATAGGGATATCAGACAG |
| 3 | P*_recF_*ΔL1 Rev | GACGGTACCCAAAACGGGAGATTTTGAAT |
| 4 | P*_recF_*ΔL 2Fwd | GACGAGCTCTATGGGTAAGAGGACTGTTTAGCA |
| 5 | P*_recO_*Fwd | GACGAGCTCACGAGAAAGAGTGGGTATA |
| 6 | P*_recO_* Rev | GACGGTACCATTAGTCATTGGTCATTAGTC |
| 7 | P*_recO_*ΔH Rev | GACGGTACCGTTAGTGAAGACAAAGGA |
| 8 | P_recO_ΔHL1 Rev | GACGGTACCGGACAAATGACTTAGGAC |
| 9 | P_recO_mutL2 Fwd | GACGAGCTCGACTTAATATATCAGGGTGCGACTAGA |
| 10 | P*_recR_*Fwd | GACGAGCTCGGATTGAGCGATAGCTTTTCAC |
| 11 | P*_recR_* Rev | GACGGTACCCCGTGGTCTTGTCTCCAGAA |
| 12 | P*_recR_*ΔL1 Rev | GACGGTACCACTATGATGGCATTTTTACACG |
| 13 | P*_recR_*ΔL2 Fwd | GACGAGCTCCTCAAACGTTAGCCTACTCAGTT |
| 14 | *ntcA* Fwd | GGCCATATGATCGTGACACAAGAT |
| 15 | *ntcA* Rev | GCGGATCCTTAAGTGAACTGTCTG |

*Underlined sequence represents *Sac*I and *Kpn*I restriction sites in forward and reverse primer sequences respectively for primers 1-13 and *Nde*I and *Bam*HI for primers 14 and 15 respectively.

**Supplementary Table S1B: List of DNA fragments cloned for promoter analysis**

| **Sr No** | **DNA Fragment Name** | **Primer Pairs used** | **Length**  **(bp)** |
| --- | --- | --- | --- |
| 1 | P*_recF_* | P_recF_Fwd & P*_recF_* Rev | 300 |
| 2 | P*_recF_*ΔL1 | P*_recF_*Fwd & P*_recF_*ΔL1 Rev | 250 |
| 3 | P*_recF_*ΔL2 | P*_recF_*ΔL2 Fwd & P*_recF_* Rev | 108 |
| 4 | P*_recO_* | P*_recO_*Fwd & P*_recO_* Rev | 235 |
| 5 | P*_recO_*ΔH | P*_recO_*Fwd & P*_recO_* ΔH Rev | 193 |
| 6 | P*_recO_*ΔHL1 | P*_recO_*Fwd & P*_recO_* ΔHL Rev | 178 |
| 7 | P*_recO_*mutL2 | P_recO_mutL2 Fwd & P*_recO_* Rev | 186 |
| 8 | P*_recR_* | P*_recR_*Fwd & P*_recR_* Rev | 174 |
| 9 | P*_recR_*ΔL1 | P*_recR_*Fwd & P*_recR_*ΔL1 Rev | 141 |
| 10 | P*_recR_*ΔL 2 | P*_recR_*ΔL Fwd & P*_recR_* Rev | 106 |
| 11 | *ntcA* | *ntcA* Fwd & *ntcA* Rev | 672 |

**Supplementary Table S1C: List of primer pairs used for qRT-PCR analysis**

| **Sr No** | **Primer Name** | **Primer Sequence** | **Gene amplified (bp)** |
| --- | --- | --- | --- |
| 1 | RecF Fwd (RT) | CAACGAGCCGTTCCCGAAAT | *recF* (150) |
| 2 | RecF Rev (RT) | TTCTGCGAGTTTGAGAGCTAAT |  |
| 3 | RecO Fwd (RT) | GGTAGGTTTTAGCATTGCGT | *recO* (133) |
| 4 | RecO Rev (RT) | TACAACTGTTTGGTAAGCAG |  |
| 5 | RecR Fwd (RT) | ACCCGCGAATTCAAAGGTAAG | *recR* (156) |
| 6 | RecR Rev (RT) | TTCGACGCTGGGACTAATTG |  |
| 7 | 16S Fwd (RT) | CAAGTCAGCATGCCCCTTACG | 16*S r*DNA (150) |
| 8 | 16S Rev (RT) | GCGATTCCTCCTTCACGCAG |  |

**Supplementary Table S2: List of Plasmids used in this study**

| **Sr No** | **Plasmids** | **Characteristics** | **Source/Reference** |
| --- | --- | --- | --- |
| 1 | pAM1956 | Kan^r^, promoter-less vector with *gfpmut2* reporter gene | Yoon and Golden 1998 |
| 2 | pAM-P*_recF_* | 0.3 kb P*_recF_* fragment cloned in pAM 1956 at *Sac*I-*Kpn*I restriction site | This study |
| 3 | pAM-P*_recF_*ΔL1 | 0.25 kb P*_recF_*ΔL1 fragment cloned in pAM 1956 at *Sac*I-*Kpn*I restriction site | This study |
| 4 | pAM-P*_recF_*ΔL2 | 0.108 kb P*_recF_*ΔL2 fragment cloned in pAM 1956 at *Sac*I-*Kpn*I restriction site | This study |
| 5 | pAM-P*_recO_* | 0.235 kb P*_recO_* fragment cloned in pAM 1956 at *Sac*I-*Kpn*I restriction site | This study |
| 6 | pAM-P*_recO_*ΔH | 0.193 kb P*_recO_*ΔH fragment cloned in pAM 1956 at *Sac*I-*Kpn*I restriction site | This study |
| 7 | pAM-P*_recO_*ΔHL1 | 0.178 kb P*_recO_*ΔHL1 fragment cloned in pAM 1956 at *Sac*I-*Kpn*I restriction site | This study |
| 8 | pAM-P*_recO_*mutL2 | 0.186 kb P*_recO_*mutL2 fragment cloned in pAM 1956 at *Sac*I-*Kpn*I restriction site | This study |
| 9 | pAM-P*_recR_* | 0.174 kb P*_recR_* fragment cloned in pAM 1956 at *Sac*I-*Kpn*I restriction site | This study |
| 10 | pAM-P*_recR_*ΔL1 | 0.141 kb P*_recR_*ΔL1 fragment cloned in pAM 1956 at *Sac*I-*Kpn*I restriction site | This study |
| 11 | pAM-P*_recR_* ΔL2 | 0.106 kb P*_recR_* ΔL2 fragment cloned in pAM 1956 at *Sac*I-*Kpn*I restriction site | This study |
| 12 | pET16b | Cb^r^, Expression vector with N-terminal His-tag | Novagen |
| 13 | pET*lexA* | Cb^r^, 606 bp *lexA* gene cloned in pET16b at *Nde*I, *Bam*HI restriction sites | Kumar et al 2015 |
| 14 | pET*ntcA* | Cb^r^, 672 bp *ntcA* gene (*alr4392i)* cloned in pET16b at *Nde*I, *Bam*HI restriction sites | This study |

**Supplementary Table S3: List of strains used**

| **Bacterial Strains** | **Characteristics** | **Source/Reference** |
| --- | --- | --- |
| ***E. coli* strains** | | |
| DH5α | F^-^ *recA*41 *endA*1 *gyrA*96 *thi*-1 *hsdR*17 (rk^-^ mk^-^) *supE*44 *relAλ* Δ*lacU*169 | Lab Collection |
| HB101 | F^-^ mc^r^ Bm^r^ rhsdS20(rB^-^mB^-^) *recA*13 *leu* B6ara-14 *pro*A2 *lac*Y1 *gal*K2 *xyl*-5 *mtl*-1 *rpsL*20 (Sm^R^) *gln*V44 λ^-^ | Lab Collection |
| Ec(pAM-P*_recF_*) | Kan^r^, DH5α/HB101 strain harbouring the plasmid pAM-P*_recF_* | This Study |
| Ec(pAM-P*_recF_*ΔL1) | Kan^r^, DH5α/HB101 strain harbouring the plasmid pAM-P*_recF_*ΔL1 | This Study |
| Ec(pAM-P*_recF_*ΔL2) | Kan^r^, DH5α/HB101 strain harbouring the plasmid pAM-P*_recF_*ΔL2 | This Study |
| Ec(pAM-P*_recO_*) | Kan^r^, DH5α/HB101 strain harbouring the plasmid pAM-P*_recO_* | This Study |
| Ec(pAM-P*_recO_*ΔH) | Kan^r^, DH5α/HB101 strain harbouring the plasmid pAM-P*_recO_*ΔH | This Study |
| Ec(pAM-P*_recO_*ΔHL1) | Kan^r^, DH5α/HB101 strain harbouring the plasmid pAM-P*_recO_*ΔHL1 | This Study |
| Ec(pAM-P*_recO_*mutL2) | Kan^r^, DH5α/HB101 strain harbouring the plasmid pAM-P*_recO_*mutL | This Study |
| Ec(pAM-P*_recR_*) | Kan^r^, DH5α/HB101 strain harbouring the plasmid pAM-P*_recR_* | This Study |
| Ec(pAM-P*_recR_*ΔL1) | Kan^r^, DH5α/HB101 strain harbouring the plasmid pAM-P*_recR_*ΔL1 | This Study |
| Ec(pAM-P*_recR_* ΔL2) | Kan^r^, DH5α/HB101 strain harbouring the plasmid pAM-P*_recR_* ΔL2 | This Study |
| BL21(*plysS*) (DE3) | Cm^r^ F^-^*ompT*hSdSB (r_B_^-^m_B_^-^) *gal dcm*pLysS (*plysS*) (DE3) | Novagen |
| BL21(*plysS*) (pET16b) | Cm^r^, Cb^r^, *E. coli* BL-21 cells harbouring the plasmid pET16b | (Kumar et al., 2018) |
| BL21 (pET16b + pAM-P_x_) | Cm^r^, Cb^r^, BL21(p*lysS*) cells having pET*lexA* plasmid, co-transformed with Kan^r^ pAM-P_x_; where P_x_ = P*_recF_*, P*_recF_*ΔL1, P*_recF_*ΔL2, P*_recO_*, P*_recO_*ΔH, P*_recO_*ΔHL1, P*_recO_*mutL2, P*_recR_*, P*_recR_*ΔL1, P*_recR_* ΔL2 | This Study |
| BL21(*plysS*) (pET*lexA*) | Cm^r^, Cb^r^, *E. coli* BL-21 cells harbouring the plasmid pET*lexA* | (Kumar et al., 2015) |
| BL21 (pET*lex*A) (pAM-P_x_) | Cm^r^, Cb^r^, BL21(p*lysS*) cells having pET*lexA* plasmid, co-transformed with Kan^r^ pAM-Px; where Px= P*_recF_*, P*_recF_*ΔL1, P*_recF_*ΔL2, P*_recO_*, P*_recO_*ΔH, P*_recO_*ΔHL1, P*_recO_*mutL2, P*_recR_*, P*_recR_*ΔL1, P*_recR_* ΔL2 | This Study |
| BL21(*plysS*) (pET*ntcA*) | Cm^r^, Cb^r^, *E. coli* BL-21 cells harbouring the plasmid pET*ntcA* | This Study |
| BL21 (pET*ntc*A) (pAM-P_x_) | Cm^r^, Cb^r^, BL21(p*lysS*) cells having pET*ntcA* plasmid, co-transformed with Kan^r^ pAM-Px; where Px= P*_recF_*, P*_recF_*ΔL1, P*_recF_*ΔL2, P*_recO_*, P*_recO_*ΔH, P*_recO_*ΔHL1, P*_recR_*, P*_recR_*ΔL1, P*_recR_* ΔL2 | This Study |
| ***Nostoc* strains** | | |
| *Nostoc*7120 | Wild type strain *Nostoc* (*Anabaena*) sp. strain PCC7120 | Lab Collection |
| AnpAM | Nm^r^, *Anabaena* 7120 harbouring the plasmid pAM1956 | Rajaram and Apte 2010 |
| AnP*_recF_* | *Nostoc* 7120 harbouring the plasmid pAM-P*_recF_* | This Study |
| AnP*_recF_*ΔL1 | *Nostoc* 7120 harbouring the plasmid pAM-P*_recF_*ΔL1 | This Study |
| AnP*_recF_*ΔL2 | *Nostoc* 7120 harbouring the plasmid pAM-P*_recF_*ΔL2 | This Study |
| AnP*_recO_* | *Nostoc* 7120 harbouring the plasmid pAM-P*_recO_* | This Study |
| AnP*_recO_*ΔH | *Nostoc* 7120 harbouring the plasmid pAM-P*_recO_*ΔH | This Study |
| AnP*_recO_*ΔHL1 | *Nostoc* 7120 harbouring the plasmid pAM-P*_recO_*ΔHL1 | This Study |
| AnP*_recO_*mutL2 | *Nostoc* 7120 harbouring the plasmid pAM-P*_recO_*mutL2 | This Study |
| AnP*_recR_* | *Nostoc* 7120 harbouring the plasmid pAM-P*_recR_* | This Study |
| AnP*_recR_*ΔL1 | *Nostoc* 7120 harbouring the plasmid pAM-P*_recR_*ΔL1 | This Study |
| AnP*_recR_*ΔL2 | *Nostoc* 7120 harbouring the plasmid pAM-P*_recR_*ΔL2 | This Study |

**Supplementary Table S4: Binding affinity of different promoter constructs of *recF/O/R* to *Nostoc* LexA**

| S. No. | Promoter construct | Binding affinity (nM) |
| --- | --- | --- |
| 1 | P*_recF_* | 100.6 ± 0.3 |
| 2 | P*_recF_*ΔL1 | 108.1 ± 0.4 |
| 3 | P*_recF_*ΔL2 | 88.2 ± 0.3 |
| 4 | P*_recO_* | 86.1 ± 0.4 |
| 5 | P*_recO_*ΔH | 105.6 ± 0.4 |
| 6 | P*_recO_*ΔHL1 | 98.2 ± 0.4 |
| 7 | P*_recR_* | 105.8 ± 0.5 |
| 8 | P*_recR_*ΔL1 | 70.3 ± 0.4 |
| 9 | P*_recR_*ΔL2 | 85.9 ± 0.7 |
