## Supplementary figures and images for "NtcA, LexA and heptamer repeats involved in the multifaceted regulation of DNA repair genes *recF, recO* and *recR* in the cyanobacterium *Nostoc* PCC7120"

### Supplementary Fig.S1

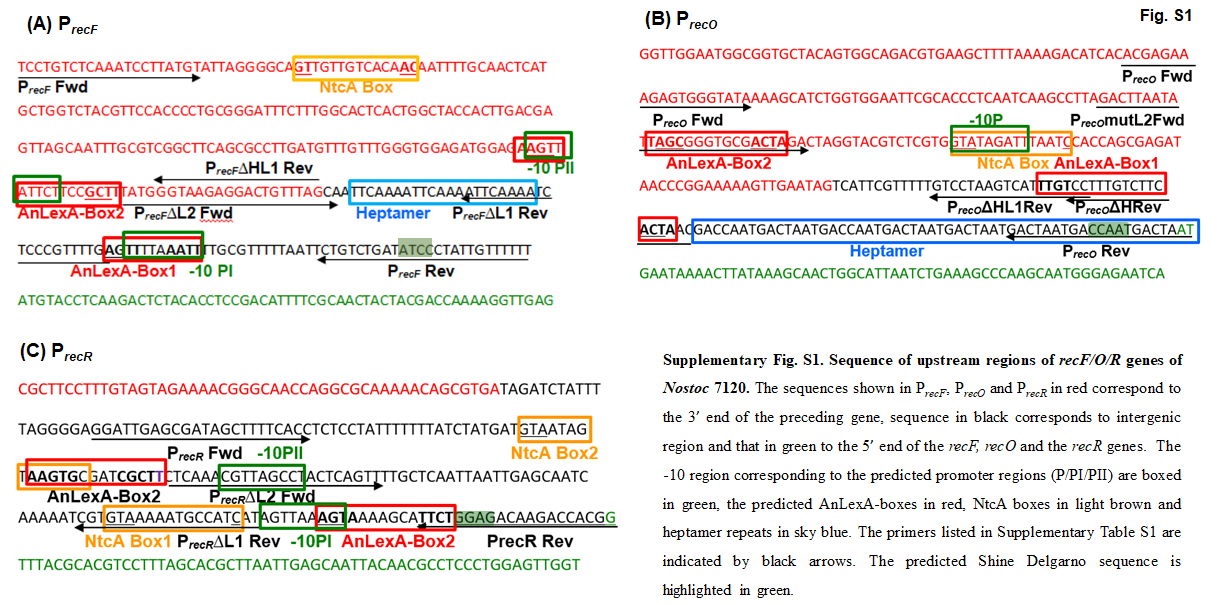

### Supplementary Fig.S2

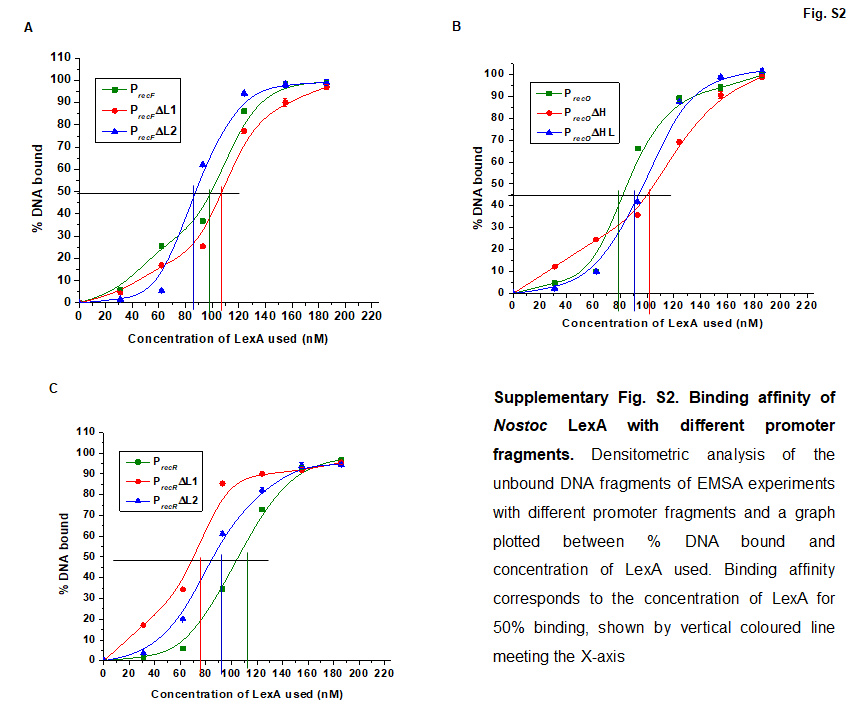

### Supplementary Fig.S3

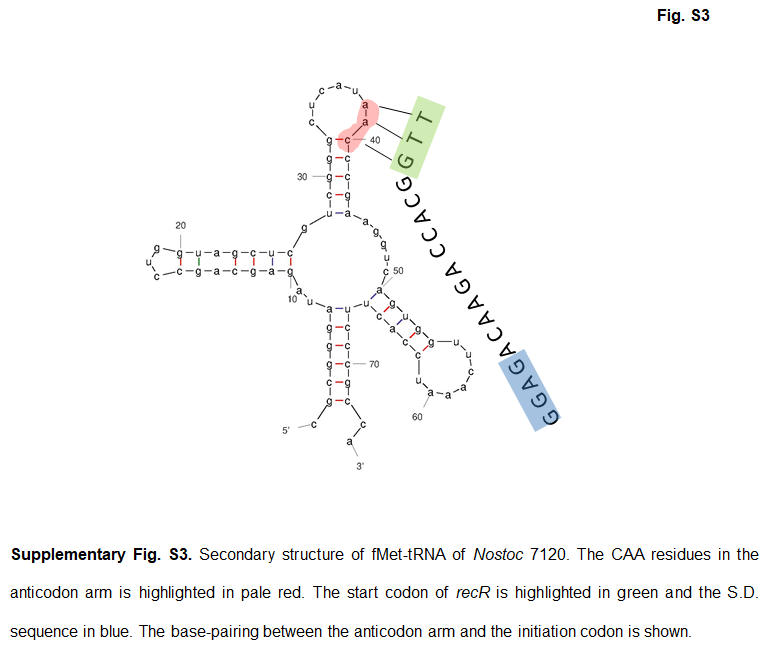
